# The lexical categorization model: A computational model of left-ventral occipito-temporal cortex activation in visual word recognition

**DOI:** 10.1101/085332

**Authors:** Benjamin Gagl, Fabio Richlan, Philipp Ludersdorfer, Jona Sassenhagen, Susanne Eisenhauer, Klara Gregorova, Christian J. Fiebach

## Abstract

To characterize the left-ventral occipito-temporal cortex (lvOT) role during reading in a quantitatively explicit and testable manner, we propose the lexical categorization model (LCM). The LCM assumes that lvOT optimizes linguistic processing by allowing fast meaning access when words are familiar and filter out orthographic strings without meaning. The LCM successfully simulates benchmark results from functional brain imaging. Empirically, using functional magnetic resonance imaging, we demonstrate that quantitative LCM simulations predict lvOT activation across three studies better than alternative models. Besides, we found that word-likeness, which is assumed as input to LCM, is represented posterior to lvOT. In contrast, a dichotomous word/non-word contrast, which is assumed as the LCM’s output, could be localized to upstream frontal brain regions. Finally, we found that training lexical categorization results in more efficient reading. Thus, we propose a ventral-visual-stream processing framework for reading involving word-likeness extraction followed by lexical categorization, before meaning extraction.

## Introduction

Reading is a crucial cultural achievement, and efficient recognition of written words is at its core. Insights have been gained into the cognitive and brain systems involved in visual word recognition (VWR), including the identification of a word-sensitive region of the left-ventral occipito-temporal cortex (lvOT). This region is often also referred to as the visual word form area (VWFA^1^); it is reliably activated by written words^2^, its structure and function are compromised in developmental reading disorders^3,4^, lvOT lesions result in severe reading deficits^5^, and electrical stimulation of this area impairs word recognition^6^. However, there is at present no agreed-upon mechanistic understanding of which process is implemented in lvOT while VWR^7^. Here, we propose a simple computational model of lvOT function during reading, i.e., the *lexical categorization model* (LCM), which integrates insights from neurocognitive and psycholinguistic research to explicitly model the response profile of lvOT to different types of orthographic stimuli.

The lvOT is part of the ventral-visual processing stream^8^. It was proposed that lvOT receives converging bottom-up visual input from both hemispheres and processes abstract representations of recurring letter sequences – including sublexical units and small words^1,9^. This proposal is in part based on the finding that lvOT is sensitive to word-<u>*similarity*</u>^10^, in the sense of decreasing lvOT activation (measured with functional magnetic resonance imaging/fMRI) with decreasing word-<u>*similarity*</u> of non-words. I.e., high activation for non-words containing letter sequences that frequently occur in real words (e.g., ‘ous’ in *mousa*) and low activation for non-words containing illegal letter combinations (e.g., *mkzsq*). Seemingly contradictory, it was reported that more <u>*familiar*</u> (i.e., more frequently occurring) words showed less lvOT activation as compared to seldom, i.e., low frequent words^11^. In sum, empirical data indicate that while word-<u>*similarity*</u> (in the sense of sub-lexical orthographic similarity) increases lvOT activation, word-<u>*familiarity*</u> (in the sense of word frequency) decreases lvOT activation. This seemingly counterintuitive set of results suggests that lvOT responds in a non-linear fashion to the ‘word-likeness’ of orthographic strings, showing greatest activity for words of intermediate word-likeness (e.g., words with low word-familiarity and non-words with high word-similarity) and least activity both for highly word-like, frequent words as well as for orthographically illegal and rarely co-occurring (‘word-un-like’) strings of letters (see also^12,13^).

The non-linear response profile of lvOT to different types of orthographic stimuli resembles the relationship between word-likeness and behavioral performance in word recognition tasks. Using a lexical decision task (categorical word/non-word decisions), Balota and Chumbley^14^ observed that lexical decisions for letter strings with intermediate levels of word-likeness were more difficult (e.g., higher error rates) than decisions to very familiar words or very ‘word-un-like’ non-words (see also^15,16^). Based on these results, they proposed that categorical recognition can be achieved for frequently occurring words and non-words that are very word-un-like (see above) exclusively on the letter string’s word-likeness. In contrast, at intermediate word-likeness levels (e.g., for rarely occurring words, words of a foreign language, or – as used in psychological experiments – orthographically legal but meaningless pseudowords), uncertainty exists concerning the lexical nature of the letter strings. Balota and Chumbley^14^ assume that this ambiguity is reduced by further analytic processing, e.g., based on sub-lexical unit (e.g., letter) processing or word-spellings.

We here propose that lvOT implements an analogous process. The core computation is a dichotomous lexical categorization primarily informed by the word-likeness of perceived orthographic strings. It is thereby yielding fast access for highly familiar words and, at the same time, filtering out uninformative non- or unknown, i.e., foreign words preventing further linguistic processing. This proposal is consistent with one of the core computational functions of the ventral-visual-stream, i.e., categorizing percepts into different categories of objects. A recent proposal described a hierarchical categorization framework of nested, spatially distinguishable cortical levels to differentiate between objects^17^. For example, animate vs. inanimate objects activate separable ventral-stream regions, while spatially dissociable sub-regions represent different lower-level features, such as faces or eyes for the animate subcategory. This object categorizing architecture is also known to categorize between words and non-orthographic objects efficiently^6,18,19^. Within the framework of hierarchically organized ventral-stream processing, it is plausible that a categorical distinction between existing words and non-words is computed at the next level of resolution (i.e., first categorizing sensory information as, orthographic stimulus, etc.; next level, within each category, categorization if it is a word) – preparing subsequent lexical, i.e., linguistic processing.

Here, we assume that the, in the lvOT implemented, lexical categorization computation precedes the retrieval of word meaning (i.e., ‘lexical access’). This computation is plausible since one saves neuronal resources efficiently enable lexical processing for familiar words. Concurrently, preventing attempts of further linguistic processing of non-words and, e.g., start a web search to identify the unknown word. Importantly, as reviewed above, lvOT activity does not linearly, but non-linearly reflect word-likeness, representing the level of uncertainty associated with the word/non-word categorization. We hypothesize that the behavioral pattern shown in lexical decision tasks reflects the processes that compute lexical status, thus providing a mechanistic explanation for the observed non-linear activation pattern of lvOT.

To characterize the lvOT’s role for word recognition in a quantitatively explicit and testable manner, we implemented this hypothesis in a simple computational model: the *lexical categorization model* (LCM). The LCM predicts the lexical categorization difficulty based on the word-likeness of the input letter string. Current conceptions of lvOT functioning in VWR are verbal-descriptive and can be divided in models that assume linear or non-linear response profiles. Linear models were the local combination detector model that suggests that lvOT activation reflects a presented letter string’s overlap with stored representations^9,10^ and the lexicon model assumes a word-frequency-based whole-word lexicon search process^11,20,21^. Non-linear response profiles were suggested besides the LCM by the engagement and effort model (E&E) and the interactive account-model (IA). The E&E accounts for a lexicon-search process and how strong a given orthographic string engages the lvOT (i.e., learned words result in higher engagement than non-words)^13^ The IA assumes predictive coding^22^ based orthographic processing, i.e., lvOT activation reflects the error related to internally generated predictions^12^. The main advantages of computational implementations are direct quantitative comparisons of alternative models, via model simulations, and direct model evaluations via correlations of simulations with empirical data.

We here evaluate the LCM against implementations from the four alternative models mentioned above and a predominant cognitive model, i.e., the Dual Route model^23^. We include simulations, reflecting the activation of the lexical (orthographic lexicon activation) and the sub-lexical route (grapheme-to-phoneme activation), both previously associated with lvOT activation (e.g.,^13,21,24^). In simulation studies, we compare all models based on nine benchmark effects reported in the literature. fMRI based evaluations allow localizing the lexical categorization computation in neuronal space. After that, models can be compared based on their simulations’ fit compared to the observed lvOT activations. Note, our primary focus is the description of lvOT responses to stimulus as opposed to task differences. Our notion here is that it is optimal to understand the activation variance determined by the stimulus characteristics, i.e., by a model, before the systematic investigation of different tasks. Our task investigation follows this logic (fMRI study 3). Finally, we investigate if training the lexical categorization process increases reading efficiency.

### LCM implementation

To implement the LCM, we first derived the word-likeness distributions of a large set of orthographic strings (Fig. 1a). We estimated word-likeness (OLD20^25^) of all German five letter words with the first letter being uppercase (i.e., nouns and noun-forms; 3,110; example: *Augen*) and the same number of pseudowords (e.g., *Augon*) and consonant strings (e.g., *Zbgtn*). The distributions of words and pseudowords overlap strongly in intermediate familiarity ranges (Fig. 1a; consistent with^14^).

**Figure 1.**
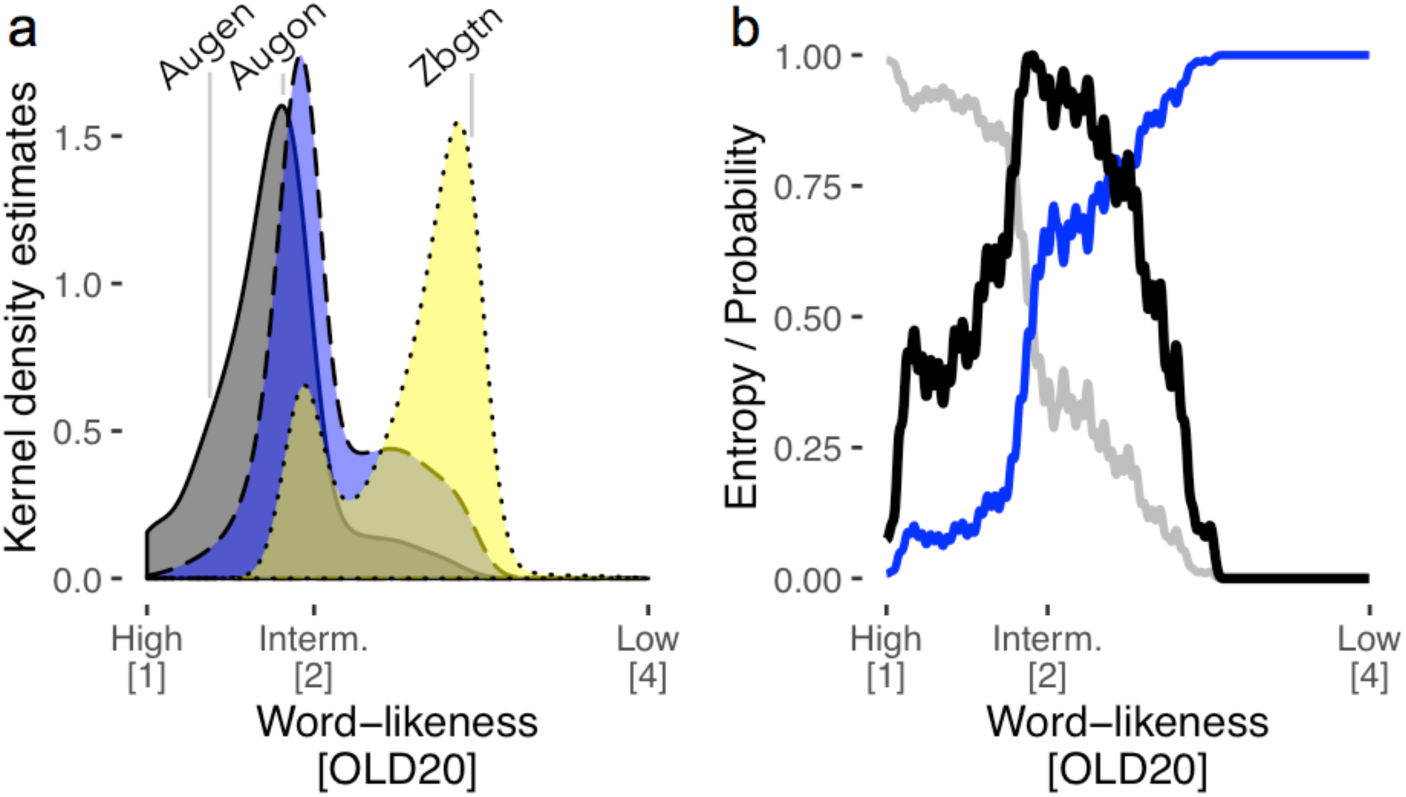
Description of the lexical categorization model (LCM). (a) Word-likeness distributions (kernel density estimates), based on the orthographic Levenshtein distance (OLD20^25^) of words (gray), pseudowords (blue), and consonant strings (yellow) including an example for each category. (b) Probability that a letter string given an OLD20 value is a word (gray line) or a non-word (blue line). The black line represents the estimated entropy (Equation 1), which combines the probabilities of being a word or non-word across all possible OLD20 values. The LCM’s central hypothesis is that this entropy function reflects lvOT activation across all possible levels of word-likeness, effectively representing the lexical categorization difficulty.

High word-likeness indicates that letter strings are highly likely words, i.e., the gray distribution showing words does not overlap with the non-word distributions. Low word-likeness (i.e., OLD20>3), in contrast, indicates that letter strings are not word-like, i.e., consonant string non-words (yellow distribution). For these strings, word-likeness estimates allow word/non-word categorizations with high degrees of certainty. This certainty is reflected in the probabilities of a string being a word or a non-word based on word-likeness (gray and blue line, respectively, in Fig. 1b). In contrast, at intermediate word-likeness levels, lexical status is ambiguous (e.g., probability of being a word/non-word is. 5; Fig. 1b). Correct word/non-word categorization is not possible based on word-likeness only. As described by Balota and Chumbly^14^, lexical categorization needs additional evidence here, e.g., spelling information, for a correct result.

Suppose the lvOT implements a lexical categorization computation, that is hard when word-likeness distributions of words and non-words overlap and easy when they do not (i.e., for frequent words or consonant cluster non-words). In that case, we expect that the non-linear response profile of lvOT is very well-described by lexical categorization difficulty. To test this assumption, we implemented the LCM using the information-theoretical concept of entropy (Fig. 1b, black line; see *Methods*).

## Results

### Evaluation A: LCM simulations

We use the LCM, implementations of four alternative neurocognitive models^1,12,13,24^ and one purely cognitive model, the Dual-Route model (DRC)^23^, to simulate the most frequently discussed published lvOT activation contrasts between different types of visually presented letter strings. We implemented the LCM simulations by transforming the word-likeness of each letter string into a lexical categorization uncertainty via the entropy function (Fig. 1b).

Figure 2a-e display LCM simulations for different types of letter strings (see *Methods*) that successfully reproduce published fMRI based lvOT activation results: pseudowords>words^13^; words>consonant strings^2^; pseudowords>words>consonant strings^26^; pseudohomophones>words and pseudohomophones=pseudowords^20^; pseudowords>words matched on multiple lexical characteristics^27^; word similarity effect: low word similarity<intermediate word similarity<high word similarity=words^10^; increasing lvOT activation with decreasing word frequency^11^ including pseudowords (note that when only words were used, the beta-weight was reduced from −0.40 to −0.17; see also^28–30^); bigram frequency effect: increasing lvOT activation with increasing bigram frequency^31^. Note that bigram frequency is also a measure previously used for estimating word-likeness and is correlated to the OLD20 parameter (e.g., see Fig. 2b in^16^). Therefore, we also present the non-linear bigram frequency effect, resolving the inconsistent findings concerning lvOT and bigram frequency previously reported^30,31^.

**Figure 2.**
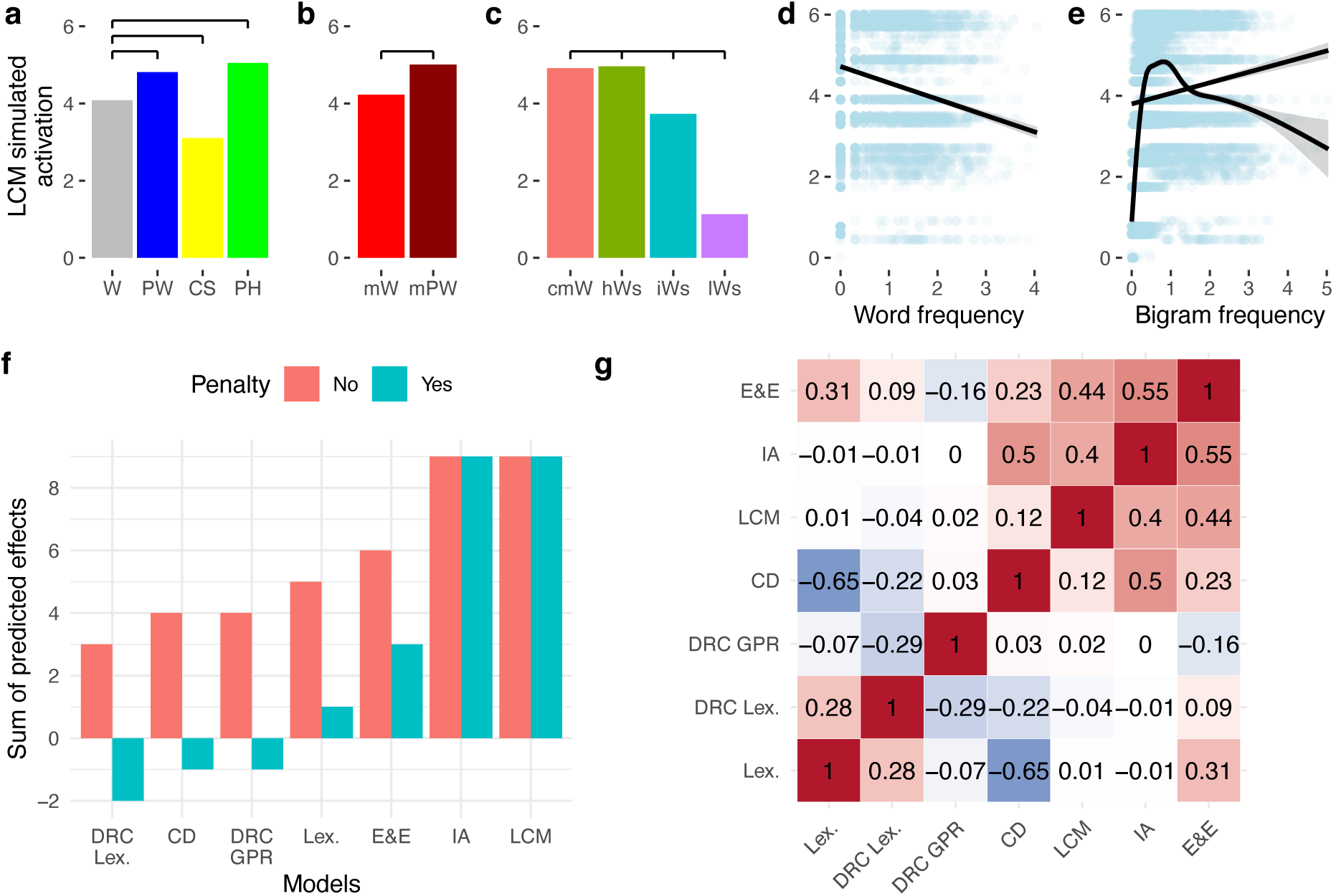
Evaluation A of the LCM based on simulations of lvOT benchmark effects from the fMRI literature and model comparisons to the lexicon model, local combination detector model (CD), effort and engagement model (E&E), the interactive account model (IA; for implementations and detailed simulations see *Supplement 1*), and the orthographic lexicon and grapheme to phoneme conversion route of the cognitive Dual-Route model (for detailed simulations see *Supplement 2*). Simulated lvOT activation (in arbitrary units: min = 0; max = 6) for all groups of letter strings is presented using bar graphs depicting their respective mean activation. In addition, horizontal black bars indicate significant differences of the simulation results between letter string categories, as derived from linear models (Bonferroni corrected). LCM simulated lvOT activation is presented, from left to right, (a) for words (W), pseudowords (PW), consonant strings (CS), pseudohomophones (PH), (b) words and pseudowords matched on number of syllables, number of Coltheart’s orthographic neighbors, frequency of the highest frequency neighbor, initial bigram frequency, final bigram frequency, and summated bigram frequency (mW, mPW), and (c) the word similarity effect comparing words (cmW: comparative matched words) to non-words with high word similarity (matched on quadrigram frequency; hWS), to non-words with intermediate word similarity (matched on bigram frequency; iWS), and, to non-words with low word similarity (lWS). In addition, (d) the word frequency effect (for all words and pseudowords as tested in^11^) and (e) the bigram frequency effect are presented as scatter plots with a linear regression line. For bigram frequency also an non-linear regression line was presented. Each dot represents one letter string, the more saturated the blue gets, the more letter strings are included. See text for more detailed description of the replicated benchmark effects including the specific stimulus sets used. (f) Qualitative model comparisons showing the sum of correctly simulated stimulus differences (orange bars) and all correct minus all incorrectly simulated effects excluding null effects. The LCM and the IA were able to correctly simulate all contrast correctly. (g) Correlation matrix of all model parameters included in the model comparison, showing that the non-linear models simulations, i.e., the LCM, E&E, and the IA model were substantially correlated (all r’s >. 4).

When we simulate these contrasts with all five neurocognitive models and the German version of the DRC, one can compare the models based on the number of correctly simulated contrasts (Fig. 2f-g; Simulations of alternative models^1,11–13^ in *Supplement 1*). Note that for the DRC, we investigated the activations of the orthographic lexicon, similar as assumed in the lexicon model^24^, and the activation of the grapheme-to-phoneme route, as suggested by^21^, reflecting letter-by-letter decoding separately. We also investigated the DRC simulated response times, but no difference to the orthographic lexicon simulations was found. This differentiation allowed us to investigate the two routes as independent hypotheses for the cognitive processes implemented in the lvOT (see *Supplement* 2 for details).

Only the LCM (Fig. 2a-e) and the IA model (Supplementary Fig. 1f) could simulate all contrasts correctly (Fig. 2f). The E&E model, also implementing a non-linear response profile, likewise performs better compared to the models assuming a linear response profile and the cognitive models. Thus, given that the most noticeable feature, which distinguishes the LCM, IA, and E&E models from the other models, pertains to the implementation of a non-linear response profile. All did reasonably well in simulating the lvOT contrasts (at least 6/9) and were also positively correlated with each other (Fig. 2g; all *r’*s>.4). Thus, we focused empirical fMRI based model comparisons on the LCM, IA, and E&E models.

### Evaluation B: fMRI measured lvOT activity

To test whether the quantitative LCM predictions capture activity in lvOT, we use three fMRI studies (cf. *Methods* for acquisition parameters and preprocessing).

#### Study 1

Participants viewed a set of letter strings covering a wide range of word-likeness (i.e., words, pseudowords, consonant strings, and strings of scrambled letters) in a block design with condition-specific blocks. Participants pressed a button whenever they detected a target (‘#####’). We generated the scrambled items by randomly replacing 90% of pixels of the monochrome word images (i.e., maximally unfamiliar; low word-likeness assumed). For all stimuli, the predicted lvOT activation was equivalent to the lexical categorization difficulty as simulated by the LCM. We used item-specific LCM simulations as a continuous predictor for the fMRI data without explicitly accounting for condition differences (event of interest: item onsets).

If the proposed categorization process is implemented in lvOT, quantitative LCM simulations should predict activation in this area. Our results support this prediction: Across the brain, only lvOT (Fig. 3a; Fig. 4; Table 1) shows a positive relationship between LCM simulations and BOLD response. Also, the detected activation cluster is consistent with previous localizations of the VWFA (including published MNI peak coordinates such as −48, −56, −16^10^). The model simulations for the three non-linear models (see simulations in Fig. 5abc) with the same stimulus material resulted in similar qualitative patterns (Consonant strings; CS< words; W < pseudowords; PW) for all three models but quantitative differences remained. For example, the difference between words and pseudowords was much smaller for the LCM compared to the IA and E&E simulations.

**Table 1.** Significant activation clusters from the fMRl evaluation with respective anatomical labels (most likely regions from the Juelich and Harvard-Oxford atlases including % overlap), cluster size (in mm3), and peak voxel coordinates (MNl space).

| Cluster | x | y | z | Mean<br>T | Max<br>T | Volume<br>[mm3] | Juelich labels | Harvard-Oxford labels |
| --- | --- | --- | --- | --- | --- | --- | --- | --- |
| **Experiment 1: LCM effect** |  |  |  |  |  |  |  |  |
| 1 | -40 | -64 | -12 | 3.1 | 4.2 | 1760 | 5.45% Visual cortex V5 L | 41.36% Left Inferior Temporal Gyrus temporooccipital part; 23.18% Left Lateral Occipital Cortex inferior division; 14.55% Left Occipital Fusiform Gyrus; 12.73% Left Temporal Occipital Fusiform Cortex; 6.36% Left Inferior Temporal Gyrus posterior division |
| **Experiment 3: LCM effect** |  |  |  |  |  |  |  |  |
| 1 | 24 | -69 | 54 | 3.6 | 4.5 | 4158 | 29.22% Inferior parietal lobule PGp R; 28.57% Superior parietal lobule 7P R; 8.44% Superior parietal lobule 7A R | 100.00% Right Lateral Occipital Cortex superior division |
| 2 | -30 | -87 | 39 | 3.8 | 5.0 | 3159 | 35.04% Superior parietal lobule 7A L; 23.93% Inferior parietal lobule PGp L; 17.09% Anterior intra-parietal sulcus hIP3 L; 8.55% Superior parietal lobule 7P L; 5.13% Anterior intra-parietal sulcus hIP1 L | 99.15% Left Lateral Occipital Cortex superior division |
| 3 | -51 | -75 | -6 | 3.8 | 5.1 | 1755 | 49.23% Visual cortex V5 L | 96.92% Left Lateral Occipital Cortex inferior division |
| 4 | -21 | -72 | 63 | 3.7 | 4.4 | 1350 | 80.00% Superior parietal lobule 7A L; 16.00% Superior parietal lobule 7P L | 100.00% Left Lateral Occipital Cortex superior division |
| **Experiment 2: Word-likeness effect** |  |  |  |  |  |  |  |  |
| 1 | -34 | -90 | -10 | 3.8 | 4.7 | 792 | 65.66% Visual cortex V4 L; 28.28% Visual cortex V3V L | 72.73% Left Lateral Occipital Cortex inferior division; 25.25% Left Occipital Pole |
| **Experiment 3: Word-likeness effect** |  |  |  |  |  |  |  |  |
| 1 | -24 | -69 | 36 | 3.9 | 5.0 | 2511 | 35.48% Anterior intra-parietal sulcus hIP3 L; 34.41% Superior parietal lobule 7A L; 17.20% Anterior intra-parietal sulcus hIP1 L | 100.00% Left Lateral Occipital Cortex superior division |
| 2 | -57 | -3 | 48 | 3.7 | 4.6 | 1134 | 52.38% Premotor cortex BA6 L; 14.29% Primary somatosensory cortex BA1 L | 88.10% Left Precentral Gyrus; 9.52% Left Postcentral Gyrus |
| 3 | -36 | -90 | 21 | 3.7 | 4.6 | 1026 | 36.84% Inferior parietal lobule PGp L; 21.05% Visual cortex V3V L; 21.05% Visual cortex V2 BA18 L; 13.16% Visual cortex V1 BA17 L | 73.68% Left Occipital Pole; 26.32% Left Lateral Occipital Cortex superior division |
| 4 | 33 | -96 | 6 | 3.6 | 4.5 | 864 | 53.12% Visual cortex V3V R; 40.62% Visual cortex V2 BA18 R | 100.00% Right Occipital Pole |
| **Experiment 2: Lexicality effect** |  |  |  |  |  |  |  |  |
| 1 | -36 | 34 | -18 | 4.0 | 6.6 | 10704 | 54.11% Broca's area BA45 L; 14.35% Broca's area BA44 L | 37.07% Left Frontal Orbital Cortex; 34.68% Left Inferior Frontal Gyrus pars triangularis; 16.52% Left Inferior Frontal Gyrus pars opercularis; 5.16% Left Frontal Pole |
| 2 | -6 | 52 | 28 | 3.8 | 5.5 | 3336 |  | 46.04% Left Superior Frontal Gyrus; 37.17% Left Frontal Pole; 5.76% Right Superior Frontal Gyrus; 5.52% Left Paracingulate Gyrus; 5.28% Right Frontal Pole |
| 3 | 56 | 32 | 10 | 3.5 | 4.3 | 1304 | 85.89% Broca's area BA45 R | 60.74% Right Frontal Pole; 24.54% Right Inferior Frontal Gyrus pars triangularis; 14.72% Right Frontal Orbital Cortex |
| 4 | -40 | -36 | -18 | 3.8 | 5.2 | 976 |  | 80.33% Left Temporal Fusiform Cortex posterior division; 18.03% Left Temporal Occipital Fusiform Cortex |
| 5 | 60 | -34 | -2 | 3.5 | 4.5 | 904 | 24.78% Insula Id1 R | 61.95% Right Middle Temporal Gyrus posterior division; 22.12% Right Middle Temporal Gyrus temporooccipital part; 15.93% Right Superior Temporal Gyrus posterior division |
| 6 | 44 | 10 | 28 | 3.5 | 4.1 | 784 | 70.41% Broca's area BA44 R | 43.88% Right Precentral Gyrus; 40.82% Right Inferior Frontal Gyrus pars opercularis; 15.31% Right Middle Frontal Gyrus |
| **Experiment 3: Lexicality effect** |  |  |  |  |  |  |  |  |
| 1 | -6 | -84 | 3 | 3.6 | 4.8 | 15687 | 23.92% Visual cortex V1 BA17 R; 21.00% Visual cortex V2 BA18 L; 19.28% Visual cortex V1 BA17 L; 7.23% Visual cortex V3V L; 6.88% Visual cortex V2 BA18 R | 33.39% Left Occipital Pole; 11.02% Right Occipital Pole; 11.02% Left Lingual Gyrus; 10.50% Right Intracalcarine Cortex; 9.12% Left Occipital Fusiform Gyrus; 8.95% Right Occipital Fusiform Gyrus; 6.37% Left Intracalcarine Cortex; 6.02% Right Lingual Gyrus |
| 2 | -57 | 12 | -9 | 3.7 | 5.3 | 9855 | 53.42% Broca's area BA45 L; 30.14% Broca's area BA44 L | 31.51% Left Inferior Frontal Gyrus pars triangularis; 20.27% Left Inferior Frontal Gyrus pars opercularis; 16.99% Left Middle Frontal Gyrus; 15.34% Left Frontal Orbital Cortex; 6.85% Left Frontal Operculum Cortex; 6.03% Left Temporal Pole |
| 3 | -48 | -72 | 21 | 3.7 | 5.4 | 7587 | 30.25% Inferior parietal lobule PGp L; 15.66% Inferior parietal lobule PFm L; 15.30% Inferior parietal lobule Pga L; 5.69% Inferior parietal lobule PF L | 33.81% Left Lateral Occipital Cortex superior division; 24.91% Left Angular Gyrus; 18.15% Left Middle Temporal Gyrus temporooccipital part; 8.54% Left Supramarginal Gyrus posterior division; 5.34% Left Lateral Occipital Cortex inferior division |
| 4 | 48 | -39 | 9 | 3.5 | 4.4 | 2943 | 31.19% Inferior parietal lobule PFm R; 20.18% Inferior parietal lobule Pga R; 8.26% Inferior parietal lobule PF R | 66.06% Right Supramarginal Gyrus posterior division; 19.27% Right Middle Temporal Gyrus temporooccipital part; 7.34% Right Superior Temporal Gyrus posterior division |
| 5 | -54 | -18 | -3 | 3.7 | 4.7 | 1593 | 8.47% Primary auditory cortex TE1.2 L | 42.37% Left Superior Temporal Gyrus anterior division; 30.51% Left Superior Temporal Gyrus posterior division; 18.64% Left Middle Temporal Gyrus anterior division; 8.47% Left Middle Temporal Gyrus posterior division |
| 6 | 36 | 33 | 3 | 3.7 | 5.1 | 1539 | 66.67% Broca's area BA45 R | 57.89% Right Inferior Frontal Gyrus pars triangularis; 24.56% Right Frontal Orbital Cortex; 17.54% Right Frontal Operculum Cortex |
| 7 | 21 | -99 | 18 | 3.6 | 4.4 | 783 | 86.21% Visual cortex V2 BA18 R; 10.34% Visual cortex V1 BA17 R | 100.00% Right Occipital Pole |

Note: Cluster-level FWE-corrected at $p < .05$ , peak-level uncorrected at $p < .001$ . All clusters with a volume smaller than 100 mm3 (i.e., only lexicality clusters) and all labels for white matter regions (i.e., from the Juelich atlas) were omitted.

**Figure 3.**
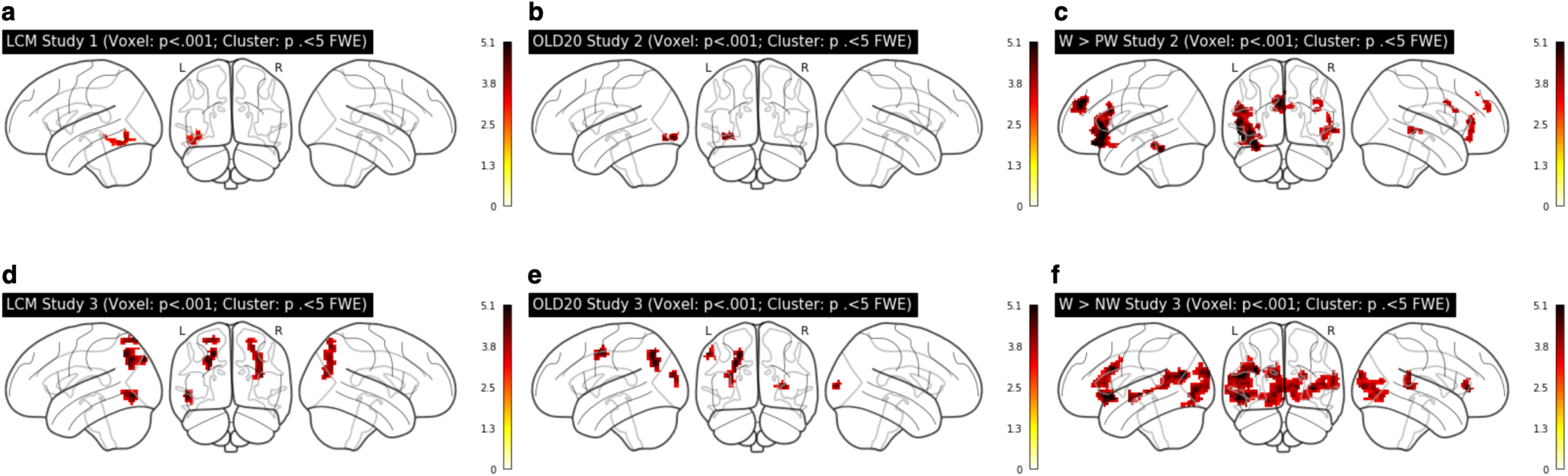
Evaluation B: fMRI whole brain analysis. Significant activation clusters using LCM simulations as a single predictor (a) in Study 1 and (d) in Study 3. Significant activation clusters using word-likeness represented by the OLD20 as a single predictor (b) in Study 2 and (e) Study 3. Words larger than non-words contrast (c) in Study 2 and (f) Study 3 (for OLD20 and word > non-word contrasts of Study 1 see Supplement 3). Thresholds for all whole brain analyses: voxel level: *p* <. 001 uncorrected; cluster level: *p* <. 05 family-wise error corrected.

**Figure 4.**
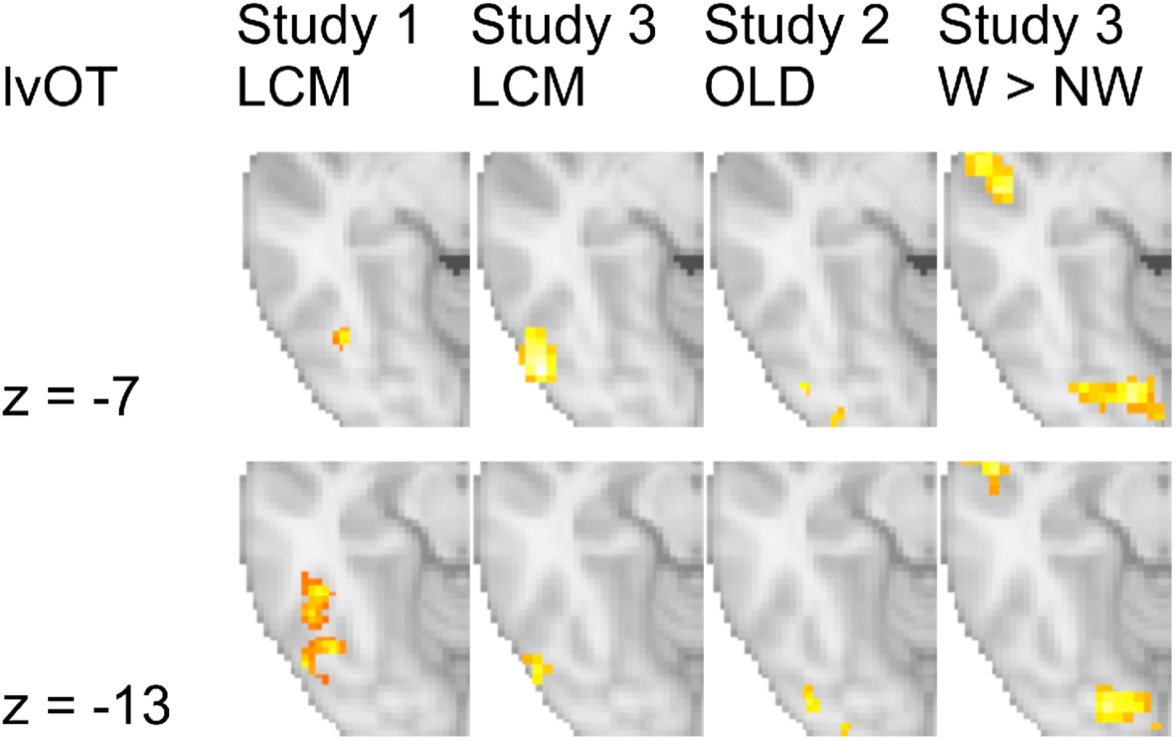
Significant LCM activation clusters in the lvOT across studies show considerable overlap, but no overlap to word-likeness and lexicality contrasts.

**Figure 5.**
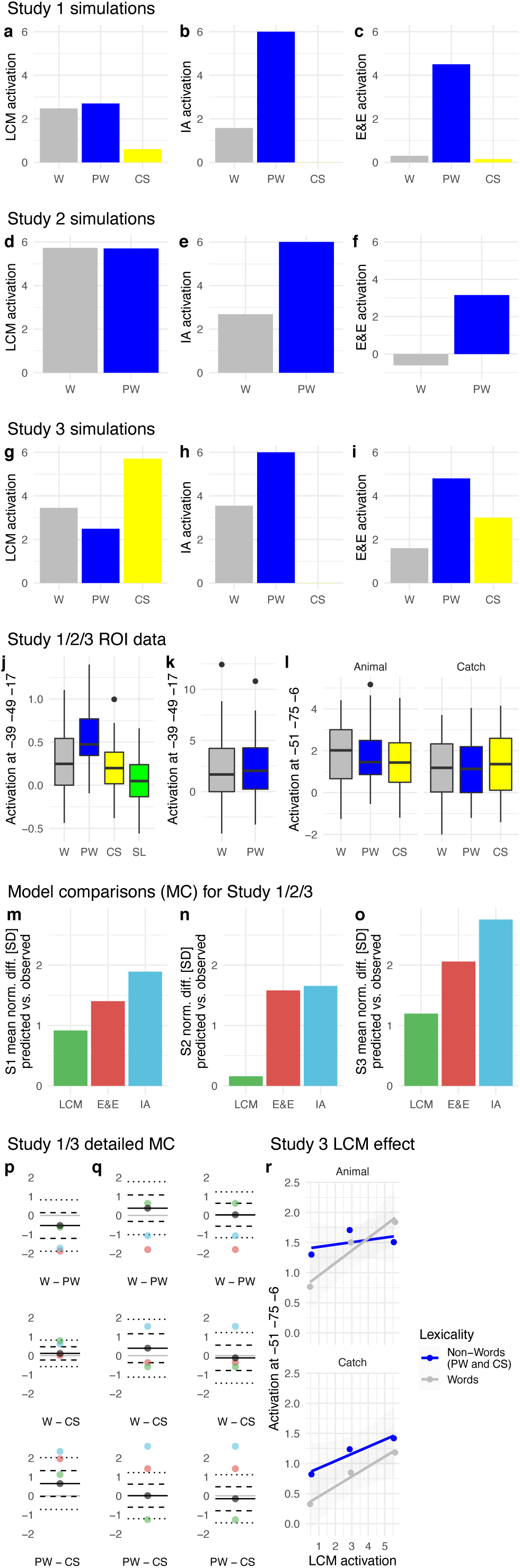
Simulations for Words (W), pseudowords (PW), and consonant strings (CS; all medians) used in Study 1 from (a) the lexical categorization model (LCM), (b) the interactive account model (IA), and (c) the effort and engagement model (E&E). (d) LCM, (e) IA, and (f) E&E simulations for Study 2. (g) LCM, (h) IA, and (i) E&E simulations for Study 3. Empirically observed lvOT activation for W, PW, CS, and scrambled letters (SL), extracted from the same peak voxel region, i.e., defined in Study 1, of interest (ROI) of (j) Study 1, and (k) Study 2. (l) For the animal decision and catch trial detection task of Study 3 the ROI data was extracted from the peak voxel of Study 3. We present the percent signal change, in arbitrary units, for each condition, including the variance across participants (horizontal line represents the median; box +− 1 standard deviation; whiskers +− 2 standard deviations). Besides, we present model comparisons for the LCM, E&E, and IA models of Study 1 (S1) in (m), of Study 2 (S2) in (n), and of Study 3 (S3) in (o). We show the mean difference between the model simulated contrasts and the observed contrast differences from the ROI data. We standardize these model comparisons via the standard deviations of the observed data (SD). I.e., the difference between simulated and observed contrast differences in standard deviations of the observed data. For Study 1, we summarize three contrasts, which can be inspected in detail in (p), Study 2 includes only one contrast presented in (n), and Study 3 combines six contrasts shown in detail in (q). In the single contrast figures (pq), the solid black line and dot show the mean observed difference, dashed lines +− 1 standard deviation, and dotted lines +−2 standard deviations also from the observed differences. Green dots show the LCM simulated contrast estimate, blue dots the IA contrast estimate, and red dots the E&E contrast estimates. In (q) the left panel represents the animal detection and the right panel the catch trial detection task. (r) The linear relationship between lvOT activation and LCM-simulations in study 3 separated for words and non-words and the two tasks conducted in the study.

The empirically measured activation patterns at the peak voxel of the lvOT cluster (Fig. 5j) also showed the same qualitative differences as simulated by the three models (CS<W<PW; Fig 5a-c). To implement a quantitative model comparison between the LCM, IA, and E&E model, we transformed all simulations and the peak data on a common scale (z-transformation). Then we compared every model’s contrast differences with the empirically measured contrast differences (i.e., W vs. PW, W vs. CS, PW vs. CS). As a difference measure, we used the standard deviation from the observed contrast (see Fig. 5m; Fig. 5p for all individual contrasts), allowing a quantitative comparison of the predicted vs. the observed contrasts. In sum, the LCM’s quantitative predictions predicted the data best, i.e., the lowest difference between predicted and observed (i.e., <1 SD; Fig. 5m). In detail, for the W vs. PW and PW vs. CS, the LCM predictions were best. Only for the W vs. CS contrast, the E&E model was more accurate. Therefore, fMRI study 1 indicated that the LCM characterizes BOLD activation patterns of lvOT during VWR.

#### Study 2

In study one, letter strings were presented in condition-specific blocks (i.e., 16 items of the same category in succession), so that the predictability of the next item concerning word/non-word categorization was high. Potentially, since the LCM does not account for the experimental context, we speculate that the blocked design may have strategically reduced the amount of processing devoted to the lexical categorization. Therefore, in the second and third fMRI study, we implemented an event-related design, randomly intermixing stimulus categories. The second fMRI study presented word-likeness matched words and pseudowords in random order (a silent reading task with catch trials, i.e., detect the German word *Taste* -button). With this stimulus material, model simulations differed between the three models: LCM simulations for these stimuli predicted high lvOT activation for both conditions with only subtle condition differences (Words>Pseudowords; Fig. 5d). In contrast, both the IA and E&E models predicted a higher activation for pseudowords than words (Fig. 5e,f). In the whole-brain contrast investigating larger activation for pseudowords than words, only a difference in the precuneus was found (see Table 1). Also, there were no differences in the lvOT, as shown by a region of interest analysis (i.e., peak voxel from Study 1). We found positive activation levels with virtually no differences between words and pseudowords (see Fig 5k). Model comparisons based on the word, pseudoword contrast showed that the LCM predicted the observed lvOT contrast best (i.e., <1 SD; Fig. 5n).

#### Study 3

Here, we selected the presented stimuli based on LCM simulations. Our notion was to create a dataset that results in a clear distinction between the three alternative models. Qualitatively the LCM-simulated lvOT activation predicted a pattern that was inverted to the pattern predominant in the literature (PW>W>CS; Fig. 2 vs. CS>W>PW; Fig. 5g). If a lexical categorization process is implemented in the lvOT than the activation should correlate with the simulations but not necessarily follow the previously described stimulus category-specific differences. For an initial finding suggesting see Graves et al.^32^ show word, pseudoword differences only when words were often used but not when they were seldom. On the contrary, the simulations of the IA and E&E models (Fig. 5hi) expected different patterns (IA: PW>W>CS; E&E: PW>CS>W) with the IA model again expecting the classic pattern shown in the literature.

Besides, we included two tasks: Animal and catch trial detection (AD and CD, respectively). Each participant conducted four runs that included either AD, CD, AD, CD or CD, AD, CD, AD, presenting the relevant words in the two tasks across participants. We assigned participants randomly to the task sequence. With this manipulation, we were able to investigate the task effect on the same set of stimuli to estimate the task’s influence on the lexical categorization process that can describe the variance between stimuli.

Similar to Study 1, we used the simulated LCM as a single predictor in a whole-brain analysis. Again we identified a correlation of lvOT activation and LCM simulations (Fig. 3d; Fig. 4; Table 1). We also found weaker correlations in the left and right parietal cortex (Fig. 3d; Table 1). Peak voxel activation of the lvOT cluster showed an LCM correlation in both words and non-words and both tasks (all t’s(34)>4.3; all p’s<. 001; Fig. 5r). In this region of interest analysis, we did not find an interaction of condition and task (all t’s<1.47; all p’s>0.14) and no significant differences between stimulus categories (all t’s<1.34; all p’s>0.17; Fig. 5l). Still, we found a task effect showing a higher activation in the animal detection task (*t*(34)=-2.2; *p*<0.03). Besides, we found that the animal detection task was harder than the catch trial detection task reflected in more errors (Animal: 4.4%; Catch: 1.5%; GLM Estimate: −0.03; SE=0.015; *t*=2.2) and longer response times (Animal: 644 ms; Catch: 520 ms; LM Estimate: −0.2; SE=0.02; *t*=9.7).

The model comparisons found that the LCM had, again, the lowest error in predicting the contrast pattern (Fig. 5o). Like in Studies 1 and 2, the E&E model simulated the lvOT pattern better than the IA model. In detail, from the six contrasts investigated, the LCM had three predicted contrasts within one standard deviation of the observed contrast difference, two that fall within two standard deviations, and one that was just outside of two (Fig. 5q). In contrast, the E&E (two <1 SD’s & four >2 SD’s) and the IA (two <2 SD’s & four > 2 SD’s) simulations resulted in much larger differences. Thus, the findings of Study 3 again replicated that the lvOT activation, while VWR, is best described by a lexical categorization computation, i.e., the LCM.

#### Word-likeness and Lexicality effects

Besides the LCM based analyses, we found correlations with word-likeness (i.e., represented by OLD20) in all three studies. We localized the word-likeness representations predominately to areas posterior to the lvOT, i.e., to occipital and parietal cortex (Fig. 3be, Fig. 4 panel 5; Supplementary fig. 3; Table 1). Also, we found one area at the precentral gyrus. A dichotomous lexicality effect (W>PW), was found most consistently in brain regions upstream to lvOT in the inferior frontal cortex (Fig. 3cf; Table 1). In Study 2, we found activation in the Superior frontal gyrus and, in Study 3, we found activation in the left superior temporal gyrus and right/left occipital pole regions.

In sum, the lexical categorization computation, represented by LCM simulations, is observed in lvOT activation patterns. Across the entire brain, the LCM’s simulation predicted activity in the often-replicated word-sensitive cluster in lvOT. Also, we described the neural correlate of word-likeness in posterior regions. Furthermore, we found dichotomous W>PW effects consistently anterior to lvOT (e.g., left inferior frontal regions). These results support our LCM proposal. I.e., that lvOT computes a dichotomous lexical categorization, using word-likeness as input.

### Evaluation C: Training based evaluation

After repeatedly finding that lexical categorization is useful to describe the activation pattern of a highly relevant brain area for reading, we here implement a behavioral training paradigm. When lexical categorization is relevant to reading performance, training of the process must determine an increase in reading speed. To test this, we implemented a training procedure for German language learners that started with an incoming assessment and three ~45 min training sessions on consecutive days and ended with an outgoing assessment. The core of the LCM training was a lexical decision task, including feedback that indicated correct responses. For comparison, we included a dataset with the same stimuli (800 words and non-words) and task from a sample of typical readers.

The analysis of response times from lexical decisions showed a non-linear effect of word likeness (i.e., OLD20) for native and non-native readers (Fig. 6a). For the non-native readers, we found a significant interaction of the lexical categorization difficulty predictor and training session. This finding indicated that the lexical categorization effect increases with training (Estimate: −0.03; SE=0.01; *t*=2.4). Besides, we identified faster response times from session to session (Estimate: −0.04; SE=0.005; *t*=8.0). However, non-native readers only reached the performance of native German readers when processing consonant strings (see Fig. 6b; Equivalence test: Upper bound: *t*(123)=-1.7; *p*=0.04; Lower bound: *t*(123)=3.7; *p*<0.001). Next, we extracted the increase of the lexical categorization effect with training for each non-native reader based on the response time data from the random slope estimates from linear mixed models. This individualized estimate represents how strong the lexical categorization effect increased with training and allows us to estimate if the training effect transfers to an increase in reading speed (administered by a pre/post-training reading speed test). We estimated a regression analysis with reading speed change as the dependent variable and the effect estimates of the individual lexical categorization effects and incoming reading speed as independent variables. Note, we also included the interaction between the two parameters. We found no significant interaction (GLM Estimate: −25.8; SE=13.4; *t*=-1.9; *p*=0.058) but a significant main effect of pre-speed (GLM Estimate: −1.1; SE=0.5; *t*=-2.3; *p*=0.022) and the lexical categorization training effect (GLM Estimate: 416.7; SE=205.4; *t*=2.0; *p*=0.046; see Fig. 6c). Overall, on average, the increase in the reading speed of the lexical categorization training was 20% (*t*(75)=4.5; *p*=0.001; Fig. 6c, right panel). Thus, this final evaluation represents evidence that lexical categorization is a core process in VWR that determines efficient reading.

**Figure 6.**
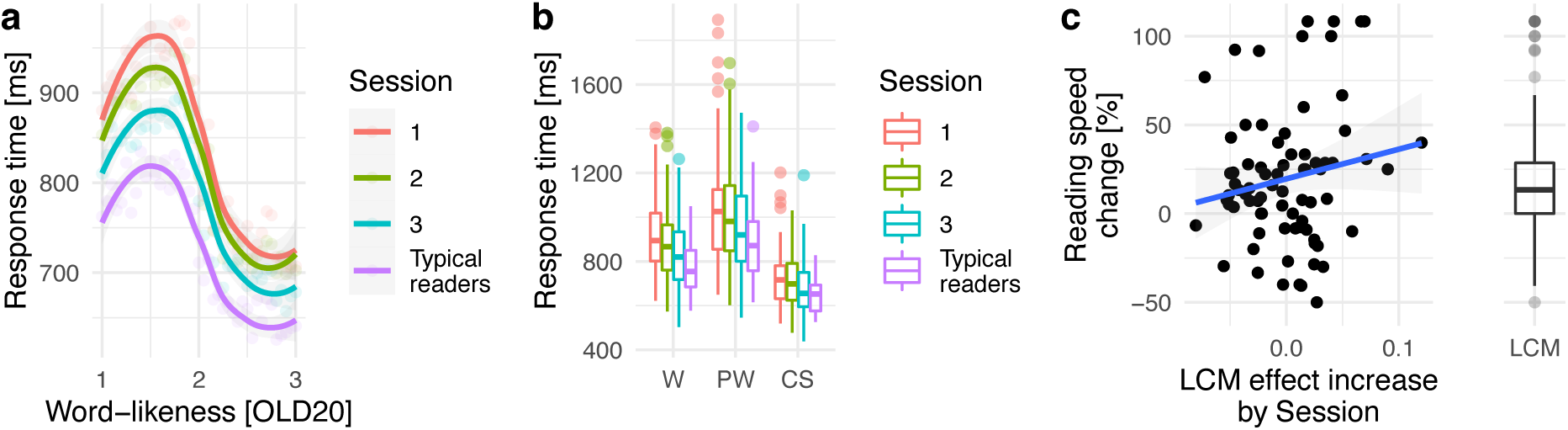
Evaluation C: Behavioral training paradigm. (a) Response times in relation to word-likeness and (b) separated for stimulus categories (W: Words; PW: Pseudowords; CS: Consonant strings) from non-native German readers while training lexical categorizations in three sessions and one group of native German readers (i.e., typical readers) doing one session of a lexical decision task with the same stimuli. (c) Reading speed change correlated with the individualized effect estimated for the LCM by training interaction. Right panel boxplot shows the overall reading speed increase. For all boxplots the horizontal line represent the median, the box +− 1 standard deviation and the whiskers +− 2 standard deviations.

## Discussion

Here, we propose that the word-sensitive cortical area in the left-ventral occipito-temporal cortex (lvOT) implements a categorization of perceived letter strings into meaningful (i.e., words) or meaningless optimizing later linguistic, i.e., lexical processing. The lexical categorization model (LCM), a computational implementation of the categorization process, is essential to our investigation. The LCM allowed understanding the intricate activation pattern of the lvOT in response to a variety of letter strings irrespective of their task context (compare^10^ and^11^). The model is capable of simulating multiple published fMRI findings and reliably predicts empirical fMRI activations in lvOT. The LCM exceeded multiple alternative models^9,12,13,23,24^, some implemented here for the first time, both in simulation and empirical studies. Finally, we found that a behavioral training procedure, based on lexical categorization, increased reading speed. Besides, we found that word-likeness, i.e., the basis of the lexical categorization, is predominantly represented in posterior occipital/temporal/parietal regions, and word-specific lexical processing predominantly is implemented in more anterior brain regions (see also^33^). These findings, thus, suggest that during reading, after initial visual processing, word-likeness is estimated in posterior occipital/parietal regions. In the following, based on word-likeness, a lexical categorization of perceived letter strings is implemented in lvOT, which precedes the extraction of word meaning, at downstream cortical sites, including, e.g., left frontal cortex (Fig. 7).

**Figure 7.**
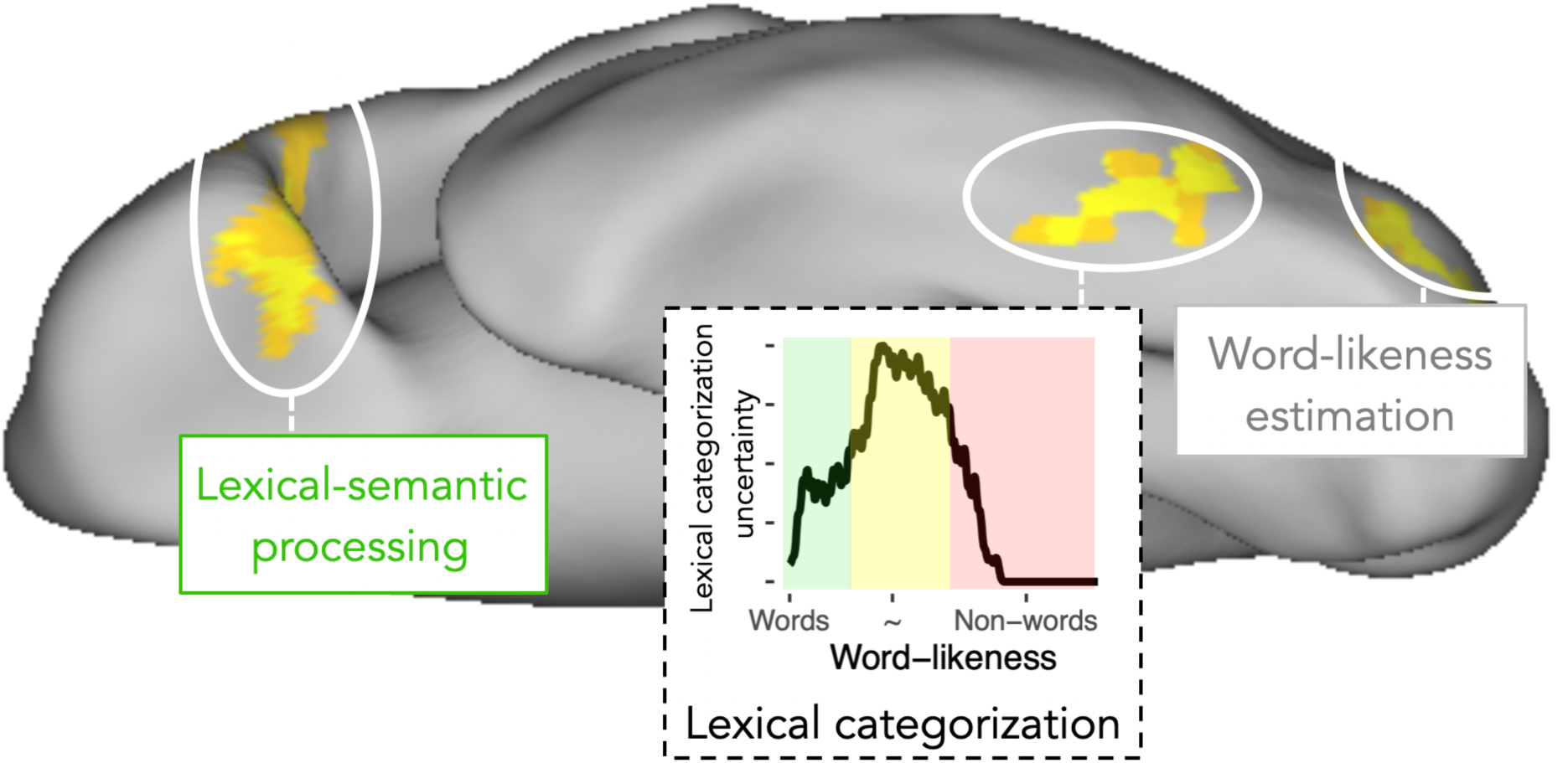
Schematic description of ventral visual stream processing during visual word recognition as assumed in the lexical categorization model, including (i) word-likeness estimations in posterior regions, (ii) lexical categorization, and (iii) the extraction of word meaning in anterior regions. For the lexical categorization process implemented in lvOT the uncertainty is presented, visualizing the assumption that higher degrees of categorization uncertainty – in areas of intermediate word-likeness – may require further elaborative processing to reach a lexical categorization.

The LCM is a quantitatively explicit extension of previous VWR models of lvOT^1,11,12^, allowing explicit comparisons to alternative models. The comparisons based on model simulations showed a clear advantage for models that considered a non-linear response profile of the lvOT, i.e., LCM, interactive account^12^, and engagement & effort model^13^. Further, fMRI data-based model comparisons showed that the LCM predicted the data best. Besides, our work differs from previous approaches by considering letter strings with a wide range of word-likeness and the utilization of an optimal word-likeness estimate (OLD20^25^; see *Supplement* 5 for LCM model simulations based on alternative word-likeness estimates). The OLD20 word-likeness estimate is theoretically meaningful, as it is a reasonable proxy for the perceptual familiarity that one acquires while becoming an efficient reader, but potentially lacks neuronal plausibility (e.g., see^16^). Also, OLD20 estimations are based on a word lexicon only (see Supplement 6 for simulations with different lexicon sizes). Still, in the entropy estimation, non-word distributions are included (for an LCM version without non-word distributions, see Supplement 1b). The latter is essential for the model’s success, which suggests that the tasks that included both words and non-words could influence our expectations of the reading material presented; thus, the non-word distributions’ inclusion becomes essential. Hence, for further investigations of this and other issues, we could use the model implementations combined with the model comparison metrics provided here as a tool for the systematic investigations of task and stimulus variance in the activation of the lvOT.

At its core, the LCM-simulated activation reflects the difficulty of the lexical categorization. It is easy to categorize letter strings as a meaningful word when the word-likeness is high. On the contrary, it is also easy to categorize a letter string as meaningless when the word-likeness is low. Both describe efficient cases of lexical categorization of letter strings, resulting in fast responses. At intermediate word-likeness levels, however, lexical categorization difficulty is high. This results from the overlapping word-likeness distributions of words and non-words – which discards word-likeness as the sole basis for lexical categorization. As a consequence, the inclusion of additional information is required for letter strings with an intermediate word-likeness and resulting in high lvOT activation and slow responses. Spelling information was brought forward as a potential source aiding categorization when it is hard, based on word-likeness only^14^. Besides, from the third fMRI study, we learned that a parietal network co-activates with lvOT, representing a potential support structure if word-likeness is not sufficient. Interestingly, the letter string-sensitive lvOT and left parietal regions’ white matter connectivity are described even before literacy acquisition^34^, and the structural connectivity in adult readers^35^. Thus, the finding of a co-activated well-connected parietal structure potentially indicates a support structure for the lexical categorization process, in case word-likeness-based differentiation is not possible. The exact specification of the information is represented in these regions’ activation has yet to be determined.

In conclusion, the LCM, which is inspired by the ventral visual processing stream models from occipital to anterior temporal and frontal regions - is a simple computationally explicit model that reliably describes lvOT activation patterns concerning a wide range of different letter strings. Empirical evaluations of the LCM support the following: Word-likeness estimation in posterior lvOT, which is then fed forward into lvOT as the basis for lexical categorization. Finally, after non-words are filtered out, and familiar words are passed through fast, higher-level cognitive processes, such as word-meaning extraction, are postulated to occur in downstream areas, including the frontal cortex. This framework (Fig. 7), including the LCM as a central cognitive process, is a significant step towards brain-based computational accounts^36^ for information processing in reading and new interventions helping slow readers.

## Methods

### LCM implementation

Given that previous work indicated a relationship between lvOT processing and word-<u>*similarity*</u>^10^ or word-*familiarity*^11^, the implementation of the LCM relies on an optimal measure of word-likeness – the mean Levenshtein distance over the 20 nearest words (i.e., the 20 words with the lowest distance; OLD20)^25^. Word-likeness distributions were derived for a larger number of orthographic strings by first estimating the Levenshtein distance, i.e., a measure of the similarity of any two strings of letters based on the number of insertions, deletions, substitutions, and/or transpositions of two adjacent letters^37^. OLD20 is considered superior to other established lexical measures^25^, such as e.g., Coltheart’s N, as it was shown to be the better predictor of behavioral word recognition performance. Also, this measure allows us to differentiate between a large range of orthographic stimuli (compare, e.g., Fig. 1a with the left panels of *Supplement* 5). For example, most non-words and a large number of words have zero orthographic neighbors. Thus, qualitatively very different letter strings (e.g., words as well as consonant strings; *Supplement* 5) have the same number of neighbors (i.e., zero). In contrast, the OLD20 can differentiate between a broader range of letter strings by describing more subtle differences in word-likeness (see above). The OLD20 is also a useful proxy of perceptual familiarity. E.g., the lexicon’s size, which is small for beginning readers and large for skilled readers, is the basis for the OLD20 estimation. When a reader has a small lexicon, the probability of finding a high number of similar words is very low since the estimation includes the 20 nearest words. With more words in the lexicon, one is more familiar with the orthography allowing a robust differentiation between words, i.e., relevant language items, and non-words, i.e., noise (see *Supplement* 6 for simulations).

OLD20 was estimated for all German five letter uppercase words (n = 3,110; extracted from N = 193,236 words of the SUBTLEX database^38^; estimated using the *old20* function of the *vwr*-package^39^ in GNU R). For each of the selected words (e.g., *Augen* -eye), we also generated a pseudoword by replacing vowels to form phonotactically and orthographically legal but meaningless letter strings (e.g., *Augon*). Pseudowords were created automatically by replacing the vowels with other vowels until the string could not be found in the SUBTLEX database anymore. Pseudowords were then revised manually based on visual inspection in order to identify illegal letter combinations. Consonant strings (i.e., orthographically illegal strings of letters; e.g., *Zbgtn*) were formed by replacing all vowels with randomly selected consonants before also computing OLD20 for each item.

Figure 1a displays the word-likeness (OLD20) distributions of these three groups of letter strings. It displays the variability of this word-likeness estimate (compare *Supplement* 5 for distributions of the same words for alternative word-likeness estimations). Some words like *Leben* (life) are more prototypical, i.e., reflected by a high word-likeness (OLD20 = 1). Others like *Fazit* (conclusion) are less prototypical, which is reflected by a lower word-likeness (OLD20 = 2.3). Note that since the OLD20 is a distance measure, higher values represent less word-like letter strings and lower values highly word-like letter strings. Some pseudowords like *Mades* (base word *Modus*/mode) are highly similar to existing words (resulting in intermediate word-likeness; OLD20 = 1.6), while most consonants strings are dissimilar to the existing words (low word-likeness for *Zbgtn*: OLD20 = 2.95). Figure 1a demonstrates that OLD20 distributions of words (grey) and pseudowords (blue) overlap strongly in intermediate familiarity ranges, consistent with the description provided by Balota and Chumbley^14^. Letter strings with the highest word-likeness are words, and, expectedly, consonant string non-words (yellow) have the lowest levels of word-likeness. Thus, the LCM rests on the assumption that for these items, one can implement lexical categorizations (word non-word decisions) with high certainty. At the same time, lexical categorization is harder at intermediate word-likeness levels.

We propose that the information-theoretical concept of entropy^40^ can describe the non-linear response profile of lvOT. We base this proposal on the assumption that lvOT implements lexical categorization to filter out perceived letter strings that are not known. Also, we assume that the uncertainty associated with this filter function in the face of overlapping distributions at intermediate levels of word-likeness reflects the difficulty (i.e., the effort) to implement the lexical categorization. Initially, entropy measures were used to determine the information value of an upcoming event in a time series. For example, in a binary categorization, if at a time point *t* the received information already allows a perfect categorization, the expected additional information value of *t*+1 is low (i.e., low entropy). In contrast, if previous information is ambiguous and does only allow a categorization around chance, the expected additional information at *t*+1 is high (i.e., high entropy) as this information might be critical for a future categorization^40^. For the current implementation, the previous information is defined by all known words – i.e., by the mental lexicon, here approximated by 3,110 five-letter words as described above. Each perceived letter string – be it a word or a non-word – can be characterized by its word-likeness, which can be quantified relative to the existing lexical knowledge, approximated in the present model by the OLD20 measure. The estimated entropy reflects the uncertainty of the lexical categorization given the word-likeness of the letter string. Note that these estimations are an approximation, based only on a subset of all possible non-words with five letters (i.e., the same number of words, pseudowords, and consonant strings: 3,110). Still, these non-words are, to some extent, related to words from the lexicon since these were the basis for constructing our non-words. Thus, non-words represent a potential source of noise added to a system based only on information from words (e.g., OLD20 estimation is related to the word items in the lexicon).

As displayed in Figure 1b, real words (grey line), most words have a high probability of being categorized as words tend to have high word-likeness (Fig. 1a). On the other hand, non-words (blue line) tend to be less word-like and are thus clearly less likely to be categorized as words. As outlined above, the lexical categorization uncertainty is particularly high at intermediate levels of word-likeness. This relationship between word-likeness and lexical categorization uncertainty is captured well by the entropy estimation, represented by the black line in Figure 1b. Entropy is low when the word-likeness estimate allows a precise categorization (as either word or non-word). Only when the word-likeness estimate indicates a considerable uncertainty concerning the lexical categorization, the entropy is high. Of note, the shape of the entropy function over word-likeness strongly resembles the non-linear response profile of lvOT discussed in the Introduction section and described behavioral performance^14,16^.

As a consequence, we here propose, as a central postulate of the LCM, that the difficulty of the lexical categorization computation (i.e., described by the entropy function; Fig. 1b) drives the neuronal activity in lvOT. Critical here is the assumption that the lexical categorization is performed based on the word-likeness of a given letter string, only. Importantly, this entropy (*Ei*) function allows to formalize the non-linear activation profile of lvOT:

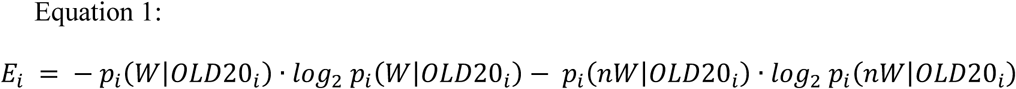

The computational implementation of the LCM consists of the entropy function (Fig. 1b, black line) derived from the probability (p_i_) of a letter string *i* being a word (W) or non-word (nW) given the specific letter string’s OLD20 (*log*_2_ indicates a logarithm on the basis of 2). *p*_*i*_(*W*|*OLD*20_*i*_) was derived by (i) taking all letter strings of a given OLD20 (e.g., at an OLD20 of 1.5 this would be 137 letter strings), (ii) identifying the words, (iii) counting them (n = 116), and (iv) calculating the probability of being a word given the OLD20 value *p*_*i*_(*W*|*OLD*20_*i*_) =. 85; *p*_*i*_(*nW*|*OLD*20_*i*_) is the inverse, i.e., 1 -. 85 =. 15).

### Participants

15, 39, and 35 healthy volunteers (age from 18 to 39) participated in Experiments 1, 2, and 3 of Evaluation B (fMRI), respectively. All had normal reading speed (reading score above 20th percentile estimated by a standardized screening; unpublished adult version of^41^), reported absence of speech difficulties, had no history of neurological diseases, and normal or corrected to normal vision. In the training study of Evaluation C, 76 healthy non-native German speaking and 48 native German speaking volunteers (age from 17–74) participated. The non-native German speaking participants were all German language learners from diverse background (Arabic, Azerbaijani, Bulgarian, Chinese, English, Farsi, French, Georgian, Indonesian, Italian, Japanese, Persian, Russian, Serbo-Croatian, Spanish, Turkish, Ukrainian, Hungarian, Urdu, and Uzbek). Also, note that six of the non-native participants became literate without the acquisition of an alphabetic script. Overall, the non-native participants had a low reading score (i.e., < percentile of 30). Selecting this group was indented as we expected that lexical categorization is well established in experienced native German readers. Participants gave their written informed consent and received student credit or financial compensation (10€/h) as an incentive for participating in the experiments. The research was approved by the ethics board of the University of Salzburg (EK-GZ: 20/2014; fMRI studies 1 and 2) and Goethe-University Frankfurt (#2015-229; fMRI study 3).

### Materials and stimulus presentation

*Evaluation A.* (i) Pseudowords>words contrast was implemented by contrasting LCM simulations of the 3,110 words and 3,110 pseudowords presented in Figure 1a. (ii) Words>consonant strings was implemented by contrasting LCM simulations of the 3,110 words and 3,110 consonant strings presented in Figure 1a. (iii) Pseudohomophones>words and (iv) pseudohomophones=pseudowords contrasts were implemented by contrasting LCM simulations of 3,110 words, 3,110 pseudowords, and 52 pseudohomophones (e.g., *Taksi*), which encompassed all 5-letter pseudohomophones presented by^21^. (v) Matched pseudowords>matched words were matched on multiple lexical characteristics, i.e., number of syllables, number of Coltheart’s orthographic neighbors, frequency of the highest frequency neighbor, initial bigram frequency, final bigram frequency, and summated bigram frequency (N = 108 vs. 108), as described in the original study reporting this benchmark effect^27^. (vi) Word similarity effect simulations are implemented with three non-word conditions including 332 letter strings with low word similarity, 4,034 letter strings with intermediate word similarity, and 220 letter strings with high word similarity, as well as 267 words. The words and non-words with high word similarity were matched on quadrigram frequency, whereas words and non-words with intermediate word similarity were matched on bigram frequency. Note that we selected the maximum possible number of items in each group to implement the match. (vii) The word frequency effect, i.e., log. counts per million, was implemented as described in the original benchmark study^11^, with N = 3,110 words and pseudowords each; the frequency of pseudowords was set to zero. (viii) Bigram frequency effect simulations are implemented including 3,110 words, pseudowords, and consonant strings each. LCM simulations of lvOT BOLD signal strength were computed as described above.

#### Evaluation B: Experiment 1

90 five-letter words, pseudowords, consonant strings, and words of scrambled letters were presented. In addition, 90 checkerboards and 16 catch trials consisting of hash marks (“#####”) were presented; to which participants responded by a button press and which were excluded from the analysis. Words and pseudowords were matched on characteristics like the OLD20, the number of syllables, and the mean bi-/tri-gram frequency (based on the SUBTLEX frequency database^38^). In addition, words, pseudowords, and consonant strings were matched on letter frequency. Stimuli were presented using Presentation software (Neurobehavioral Systems Inc., Albany, CA, USA) in black courier new font on a white background for 350 ms (1,000 ms inter-stimulus interval/ISI), in six blocks per stimulus category with 16 items each. After two blocks, a fixation cross was presented for 2 s. In addition, six rest blocks (fixation cross) were interspersed. Each block lasted for 16 s, which resulted in approximately 10 min of recording time.

#### Evaluation B: Experiment 2

60 critical five-letter words and pseudowords were presented. In addition, 120 different pseudowords that underwent a learning procedure and, thus, not analyzed here. 10 practice trials, and 30 catch trials consisting of the German word *Taste* (Button) were presented. Participants responded to catch trials by a button press; these trials were excluded from the analysis. All words and pseudowords consisted of two syllables and were matched on OLD20 and mean bigram frequency. Letter strings were presented by the Experiment Builder software (SR-Research, Ontario, Canada) for 800 ms (yellow Courier New font on gray background; ISI 2,150 ms). To facilitate estimation of the hemodynamic response, an asynchrony between the TR (2,250 ms) and the stimulus presentation was established. In addition, 60 null events (fixation cross as in the ISI) were interspersed among trials. The sequence of presentation was determined by a genetic algorithm^42^, which optimized for maximal statistical power and psychological validity. The fMRI session was divided in two runs with a duration of approximately 8 min each.

#### Evaluation B: Experiment 3

We presented 200 critical five-letter words, 100 consonant strings, and 100 pseudowords. Besides, we presented ten practice trials and 100 catch/animal trials, each consisting of the German word Taste (Button) or animal names. Participants responded to catch trials in the catch trial task or the animal names in the animal detection task by a button press; we excluded these trials from the analysis. We selected the letter strings based on simulations from the LCM, the IA, and the E&E model so that the stimulus set allowed to differentiate between the alternative models (i.e., see Fig. 5g-i). Letter strings were presented by the Presentation software (Neurobehavioral Systems Inc., Albany, CA, USA) for 800 ms (black Courier New font on gray background; ISI 2,150 ms; i.e., same as for Experiment 3). Also, 100 null events (fixation cross as in the ISI) were interspersed among trials. A genetic algorithm again determined the sequence of the presentation^42^. We divided the fMRI session into four runs with a duration of approximately 10 min each.

#### Evaluation C

We presented 1,600 5-letter letter strings in the training study, including 800 words, 400 pseudowords, and 400 consonant strings. These letter strings varied naturally on word-likeness (OLD20), word frequency, and others as we randomly drew the words and non-words from the distributions presented in Figure 1a. This procedure’s motivation is to sample from the full distribution as a critical feature of our new training, i.e., preventing artificial sets of stimuli. Thus, participants were allowed to learn based on a representative set of words. Stimulus presentation was implemented with the Experiment Builder software (SR-Research, Ontario, Canada) using mono-spaced Courier-New font, with a visual angle of approximately 0.3° per letter. The letter string presentation was randomized for each participant and each of the three training sessions. After each trial, participants got feedback if their response was correct or not. Before and after the training sessions, we assessed reading speed by a standardized screening. The latter allowed us to estimate the transfer effect of the LCM training on reading speed. Note that we combined this dataset from three studies. Note that all stimuli including lexical characteristics will be available on the Open Science Framework.

### Data acquisition and analysis

#### LCM simulations

Statistical comparisons of the simulations presented in Figure 2 were implemented with the *lm* function in R and *p*-values were Bonferroni-corrected for multiple comparisons. In total, nine benchmark effects were tested of which the contrast pseudowords>words>consonant strings^26^ was a combination of the pseudowords>words^13^ and the words>consonant strings^2^ contrast. Significant differences were marked in Figure 2 (also implemented for alternative models presented in the *Supplement* 1 and 5 with a black horizontal bar when the direction of the effect was expected from the literature and a red bar when the expected effect direction was violated. Figures 1, 2, 5 and 6 were implemented using *ggplot2* in R.

#### fMRI data; Experiments 1, 2 &3

A Siemens Magnetom TRIO 3-Tesla scanner (Siemens AG, Erlangen, Germany) equipped with a 12-channel head-coil (Experiment 1), a 32-channel head-coil (Experiment 2) or an 8-channel head-coil (Experiment 3) was used for functional and anatomical image acquisition. The BOLD signal was acquired with a T2*-weighted gradient echo-planar imaging (EPI) sequence (TR = 2250 ms; TE = 30 ms; Experiment 1&2: Flip angle = 70°, Experiment 3: Flip angle = 90°; Experiment 1&3: 64 x 64 matrix; FoV = 210 mm; Experiment 2: 86 x 86 matrix; FoV = 192 mm). Thirty-six descending axial slices with a slice thickness of 3 mm and a slice gap of 0.3 mm were acquired within the TR. In addition, for each participant a gradient echo field map (Experiment 1&2: TR = 488 ms; TE 1 = 4.49 ms; TE 2 = 6.95 ms; Experiment 3: TR = 650 ms; TE 1 = 4.89 ms; TE 2 = 7.35 ms) and a high-resolution structural scan (T1-weighted MPRAGE sequence; Experiment 1&2: 1 x 1 x 1.2 mm; Experiment 3: 1 x 1 x 1 mm) was acquired. Stimulus presentation was implemented, in Experiment 1&2, by an MR-compatible LCD screen (NordicNeuroLab, Bergen, Norway) and, in Experiment 3, by a Sanyo PLC-XP41-projector (SANYO Electric Co., Osaka City, Japan), both with a refresh rate of 60 Hz and a resolution of 1024×768 pixels.

For experiment 1 and 2 the SPM8 software (http://www.fil.ion.ucl.ac.uk/spm), running on Matlab 7.6 (Mathworks, Inc., MA, USA), was used for preprocessing and statistical analysis. Functional images were realigned, unwarped, corrected for geometric distortions by use of the FieldMap toolbox, and slice-time corrected. In Experiment 1 the high-resolution structural image was pre-processed and normalized using the VBM8 toolbox (http://dbm.neuro.uni-jena.de/vbm8). The image was segmented into gray matter, white matter and CSF, denoised, and warped into MNI space by registering it to the DARTEL template of the VBM8 toolbox using the high-dimensional DARTEL registration algorithm^43^. Based on these steps, a skull-stripped version of the structural image was created in native space. The functional images were co-registered to the skull-stripped structural image and then the parameters from the DARTEL registration were used to normalize the functional images to MNI space. In Experiment 2 the images were co-registered to the high-resolution structural image, which was normalized to the MNI T1 template image. The functional images were further resampled to isotropic 3 × 3 × 3 mm voxels in Evaluation B, Experiment 1, and 2 × 2 × 2 mm voxels in 2, Experiment 2, and smoothed with a 6 mm full width half maximum Gaussian kernel. For Experiment 3, we first set up the fMRI data in the BIDS format^44^, which allowed us to use the fMRIPrep preprocessing pipeline^45^.

For statistical analysis, in Experiment 1, 2, & 3, a two-stage mixed effects model was used. The first level is subject–specific and models stimulus onsets with a canonical hemodynamic response function and its temporal derivative. Movement parameters from the realignment step and catch trials were modeled as covariates of no interest. A high-pass filter with a cut off of 128 s was applied to the functional imaging data and an AR(1) model^46^ corrected for autocorrelation. In Experiment 3, we used the Python-based *nistats* package for statistical analysis and the *nilearn* package to create figures^47^ to model the fMRI statistics as in the SPM software. For the statistical analysis of ROI data, LMMs^48^ were calculated in R (see below).

#### Training data

Linear mixed model (LMM) analysis is a linear regression analysis that is optimized to estimate statistical models with crossed random effects for items^48^. These analyses result in effect size estimates with confidence intervals (SE) and a *t*-value. *t*-values larger than 2 are considered significant since this indicates that the effect size ±2 SE does not include zero^49^.

For the training data, we used LMMs to analyze the response time data from the LCM training task (i.e., a lexical decision with feedback). First, we excluded response times below 300 ms and above 4000 ms. Second, we log. transformed the response times to account for the ex-gaussian distribution that typically results from response times measurement. For the regression model, which we used to estimate the change of the entropy effect with training, the core term was a three-way interaction of training session, entropy, and word-likeness (i.e., OLD20). Besides, we controlled for the following covariates: word frequency, lexicality, trial index (i.e., at which position in the training session the letter string was presented), and if the response was erroneous or not. Random effects were the intercepts of letter string and participant.

For this secondary analysis, i.e., correlating individual LCM training effects with reading speed increase, we estimated the random slope of the interaction of entropy with training on the participants. With this individualized interaction estimate, we now have a predictor that can investigate if the LCM specific training effects translate to a reading speed increase. For this analysis, we estimated a linear regression model that predicted the reading speed increase in percent. As predictors, we included the pre-training reading speed and the individual LCM training estimate plus the two parameters’ interaction.

### Data and Code availability

Data and code will be published at the Open Science Framework when accepted.

## Author contributions

B.G., J.S. and C.F. conceptualized the model and wrote the manuscript. B.G. implemented the model, model simulations and alternative models. B.G., S.E., F.R. and P.L. designed, conducted, and analyzed the fMRI experiments. B.G. and K.G. designed, conducted, and analyzed the training study.

## Acknowledgements

We thank Anne Hoffmann, Jan Jürges, and Rebekka Tenderra for help with fMRI data acquisition and Benjamin Peters for help with the entropy formulation. In addition, we thank Sophia Haan for helpful comments on a previous version of the manuscript. The research leading to these results has received funding from the European Community’s Seventh Framework Programme (FP7/2013) under grant agreement n° 617891 awarded to CJF and from the European Community’s Horizon 2020 Programme (2016) under grant agreement n° 707932 awarded to BG.

## Supplementary material

**Supplementary figure 1.**
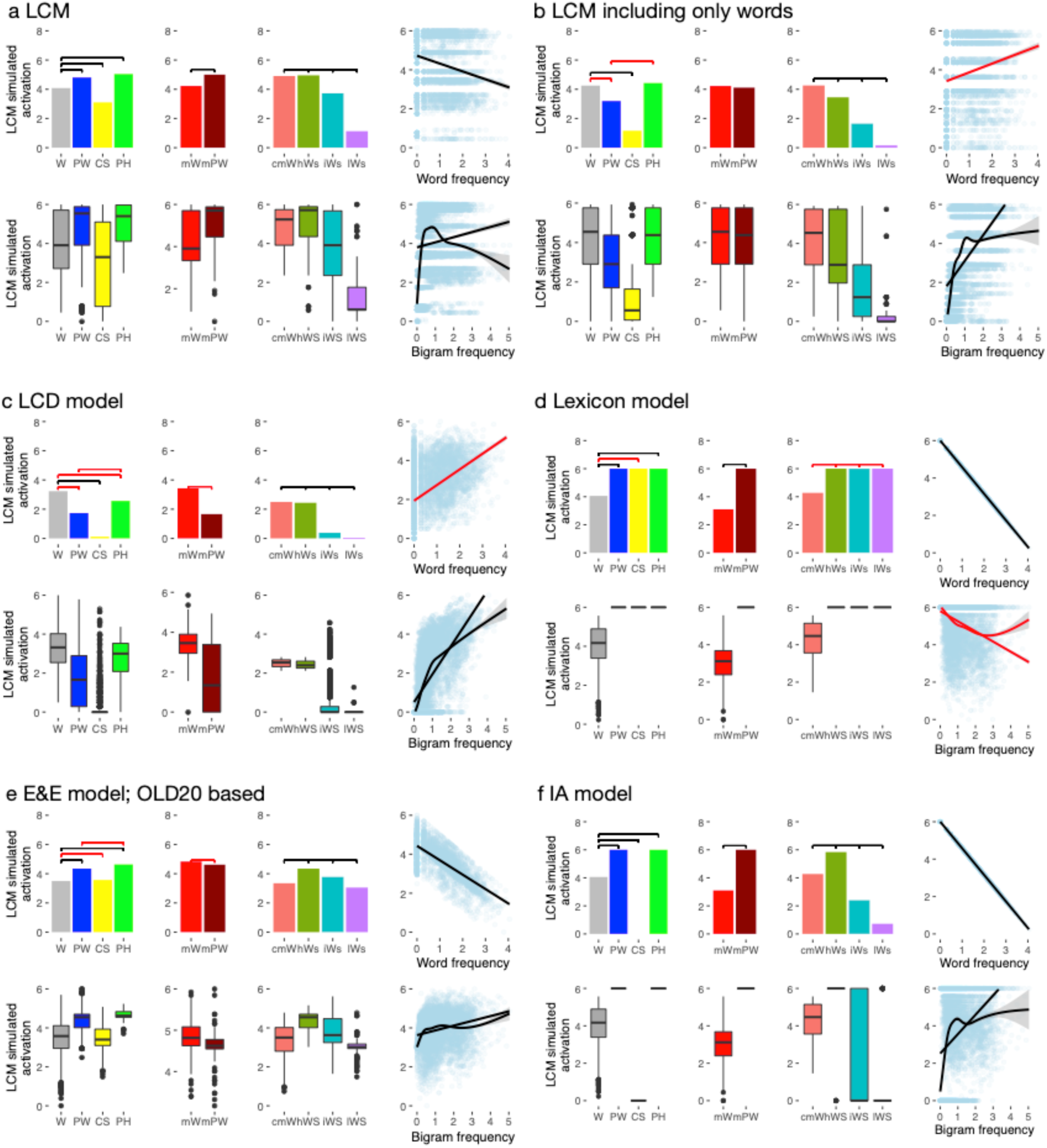
Detailed depiction of simulated lvOT activation from computational implementations of alternative models: (a) the LCM (similar as in Fig. 2, adding boxplot figures), (b) an LCM variant that included only the distributions of words, (c) the local combination detector model (LCD), (d) the lexicon model, (e) the engagement and effort model (E&E), and (f) the interactive account model (IA). Above each subplot, horizontal black bars indicate simulations that resulted in the expected condition difference between letter string categories, as derived from linear models. Red bars indicate contrasts opposite to the expected direction. In the following the details of each alternative model will be described, which allow to evaluate the LCM in contrast to alternative models of lvOT function. We implemented multiple accounts based on the verbal proposals^11,20,1,9^ and a variant of the LCM which was not described previously. The critical difference between the LCM variant and the LCM presented previously (Fig. 1) is that in the variant, we used only the word-likeness distribution of words as the basis for the entropy estimation. This implementation’s primary motivation was that it would be reasonable to test if the entropy function can be implemented based on words only. If the variant LCM is better in simulating lvOT activation contrasts, one could argue that the representations in the lvOT are, to a great extent, words. Alternatively, i.e., if the LCM model described in the main text results in more accurate simulations, one could argue that the representations in the lvOT are, to a great extent, pre-lexical.

For the implementation, we used the cumulative probability of words based on word-likeness to estimate the probability of a sting being a word/pseudoword. In other words, we took all 3110 words and estimated the probability of all words starting with the highest word-likeness level.

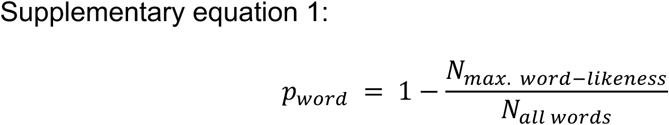

In the following, we repeated this procedure for all word likeness levels to estimate the probability for a word for each bin. In the end, the cumulated probabilities are 1.

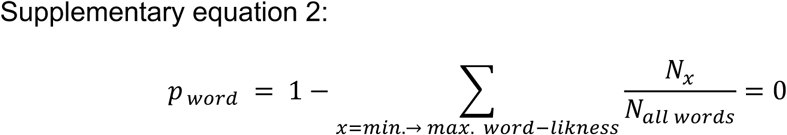

The inverse of this probability for the string being a word is again the probability of being a non-word. After that, we can use the same equations as presented in the methods to estimate the entropy, i.e., the simulated activation of the lvOT. The simulation results show that the simulations of this words-only LCM are only correct for four benchmark effects (Supplementary Fig. 1b; 4/9 effects simulated correctly). Thus, we learn here that the non-words distributions are a necessity for the LCM, suggesting that representations are, to some extent, pre-lexical in nature (see also^10^).

As an adequate test case for the LCM, we implemented four alternative models, i.e., two linear models: lexicon and LCD model, and two non-linear models, the E&E and the IA model. Here we focused on implementing the verbally described relationships between lvOT activation and the word characteristics central to the respective model. For the E&E and the IA models, we implemented more variants, but we only present those with the best fit. Overall, only the LCM and the IA were able to predict all nine benchmark contrasts. For qualitative model comparisons and model performance, see Figure 2f.

The *LCD model*^1^ postulates that lvOT stores pre-lexical and lexical multi-letter representations (up to a length of about four letters). When a letter string is presented that contains these multi-letter combinations, their neural representations are activated irrespective of the lexicality of the string. Therefore, higher activations (_-456_) are predicted for often-occurring letter combinations in contrast to rare letter combinations. As formalized in Suppl. Equation 3, this is modeled by a linear increase of 0.4 per log quadrigram frequency (of a string). The specific value for this increase function was read out from Figure 4 of^10^ (left hemisphere, y = −56;).

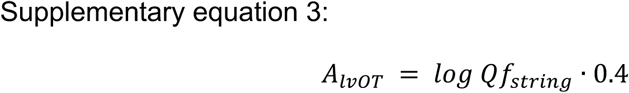

This ‘(sub-) lexical storage/activation’ model of lvOT accounts for four benchmark contrasts (Supplementary Fig. 1c). The LCD model predicts increased activation for words relative to consonant strings, as well as effects of word similarity and bigram frequency.

The *lexicon model*^24^ conceptualizes the processes implemented in lvOT as a search in an orthographic lexicon. This lexicon is arranged by word frequency; accordingly, frequent words are found fast, resulting in fast response times and low lvOT activation, whereas it takes substantially longer to identify infrequent words, resulting in prolonged response times and high activation. lvOT activation (*A*_*IvOT*_; cf. Supplementary equation 4) decreases linearly by about 1 activation level (arbitrary unit) per 1 log frequency (*f*_*string*_) unit (read out from Figure 2: Midfusiform/Posterior fusiform in^11^; *f*_*string*_: frequency of the letter string, with frequencies of pseudo- and nonwords set to zero^11^; *f*_*max*_: highest frequency in the lexicon).

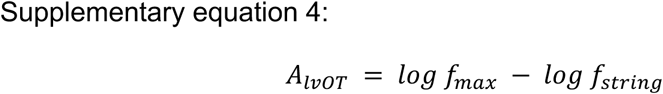

As shown in the upper panel of Supplementary Figure 1, the implementation of this model can account for five benchmark effects. Specifically, this ‘frequency ordered lexical search’ model obviously accounts for the word frequency effect but also for the lower activation for words in contrast to pseudowords irrespective if they were matched for OLD20 or not (black lines in Suppl. Fig. 1d).^13^ None of the benchmark effects including consonant strings or word similarity manipulations (i.e., bigram frequency and word similarity effect) could be captured by this model.

None of the benchmark effects explained by the lexicon model could be explained by the LCD model and vice versa. Note that the patterns of predictions generated by these two models do not depend on the numeric values chosen for the change in activation, which were read out from the results figures of the original publications. Given that these two models are simple linear transformations of the respective lexical measure, the effect directions would be similar irrespective of which specific constant number would be used to predict lvOT activation in Supplementary Equations 3 and 4. In the following we present two models implementing a non-linear relationships as the basis for the activation pattern in the lvOT in response to words.

The *engagement and effort model*^13^ combines two processes. One relates to the engagement of brain regions in visual word recognition. Engagement is high for letter strings that are likely words and low for i.e., consonant strings that violate basic rules for constituting orthographic stimuli (e.g., no vocals present). The second process is concerned with the effort one needs to recognize a word, e.g., a seldom word is more effortful to decode then a frequently used word. Here we used OLD20 to implement the engagement process, as the OLD20 parameter can differentiate between non-words. Note that we have to invert the OLD20 as it is a distance measure. Then a high value indicates a high word likeness, i.e., a high engagement of the region. To model the reduction of activation with reduced effort in processing, we subtracted word frequency from the engagement variables (i.e., both parameters were z-transformed). The result of this estimation is a non-linear response function of the lvOT with low activation for non-words with a low word-likeness, a high activation for pseudowords, and a reduced activation for words with a high frequency.

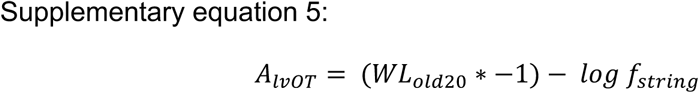

The simulations show that the E&E model could simulate six benchmark effects correctly. This finding indicates that the non-linear models are better in explaining lvOT activation in response to various letter strings.

The *interactive account* of Price and Devlin^12^ adopts a predictive coding perspective and postulates that lvOT activation reflects an interaction between top-down and bottom-up processes, in the sense of a prediction error that represents difference between, in the case of single word presentation, non-strategic general expectations derived from general word knowledge and the actual bottom-up visual orthographic input. Here we implement this by assuming bottom-up information as word frequency (i.e., often seen words are more likely expected). This reflects that bottom-up information, in standard single word presentations, is determined by predictions based on our orthographic knowledge (e.g., see^16^ for an example and more deep discussion on this issue). This is as for seldom words or non-words sensory information cannot be predicted to a high extent resulting in a large prediction error. In contrast, predictions are more precise for frequent words and therefore the prediction error is reduced. In their paper, Price and Devlin state that this in only the case for letter strings that follow the basic rules of orthography. I.e., consonant strings, that do not fall in this category therefore should not activate the lvOT. As a consequence this model also assumes a non-linear response function of the lvOT.

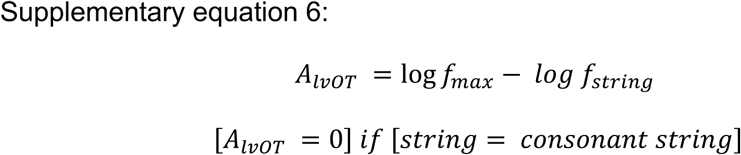

Besides the LCM, the IA model, is, the second model approach that is able to simulate all benchmark effects. This finding once more, indicates that the lvOT follows a non-linear response profile but also strongly suggests that we need experimental data to distinguish between the LCM and the IA model (see Evaluation B).

**Supplementary figure 2.**
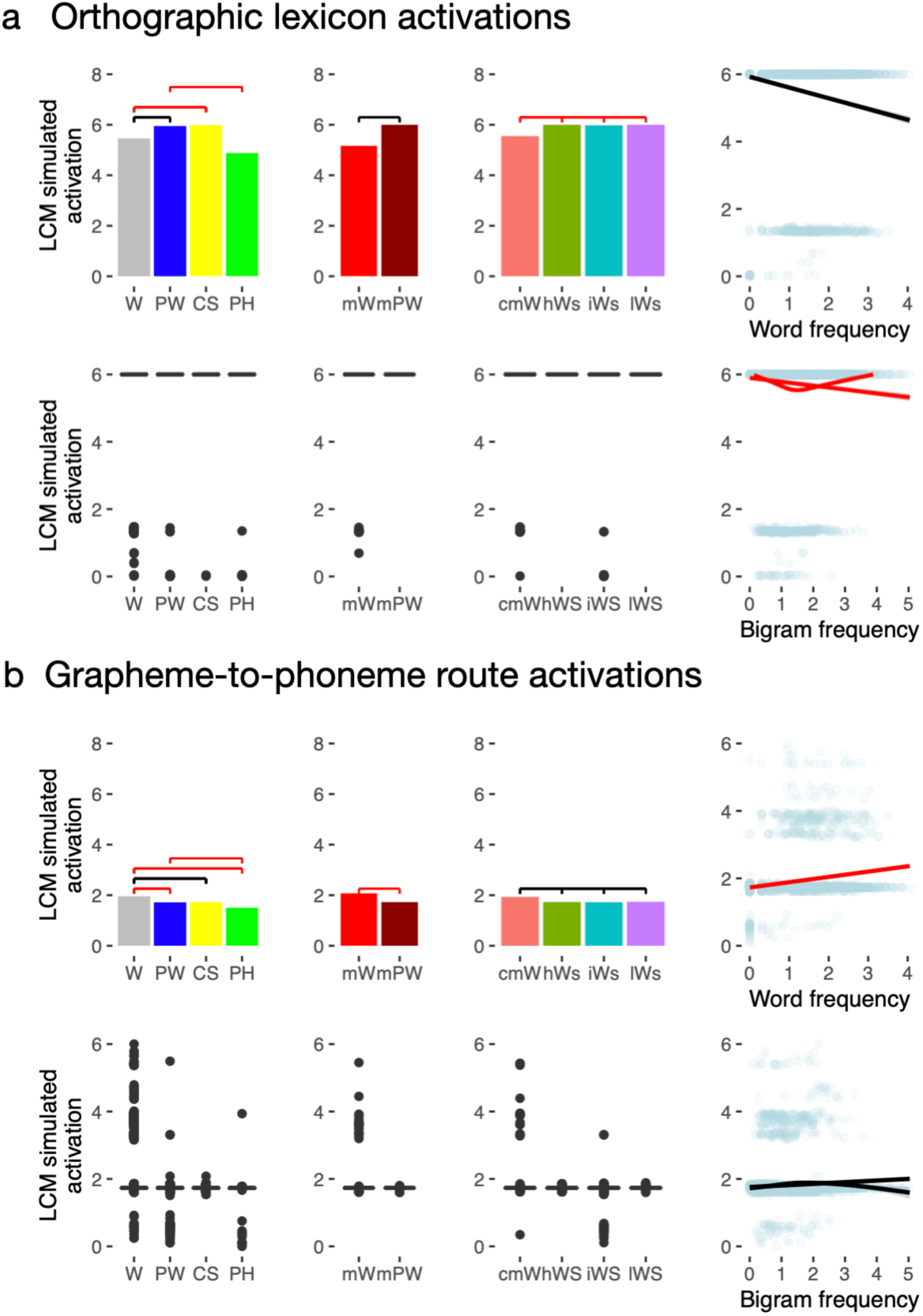
Simulations from the Dual-Route Model of reading aloud (DRC; Coltheart et al., 2001) separated for activation from (a) the orthographic lexicon and (b) the grapheme-to-phoneme conversion route of the DRC. We estimated the activation per word by the normalized correlation of model cycles and the activation level. This metric results in a low activation when the orthographic lexicon activation rises fast. The former reflecting easy access, and the latter reflecting hard access to the lexical item. We applied the same logic to the grapheme-to-phoneme route. Here a fast activation rise indicates that the letter string can be decoded fast based on the letter-by-letter decoding process. Again a fast activation rise of that route is reflected in a low correlation. In contrast, a slow rise is reflected by a high correlation, i.e., assumed high activation. Interestingly, the orthographic lexicon activation resulted in a simulated activations pattern that mimicked the lexicon model, and the grapheme-to-phoneme route the local activin detector model simulations. All but the high activation for pseudohomophones could be simulated by either the orthographic lexicon or the grapheme-to-phoneme route. Thus, one can conclude that the DRC can, when orthographic lexicon and grapheme-to-phoneme route are both implemented in lvOT, explain most of the pattern found in the lvOT by one or the other route of the model. Still, when using the reaction times from the model, i.e., the simulations using both routes, the expected contrast differences could not be simulated better than the orthographic lexicon results. Nonetheless, the simulated activation for pseudohomophones was lower than one would expect from the literature pattern. I.e., simulated activations were lower than words, pseudowords, and consonant stings in both routes.

**Supplementary figure 3.**
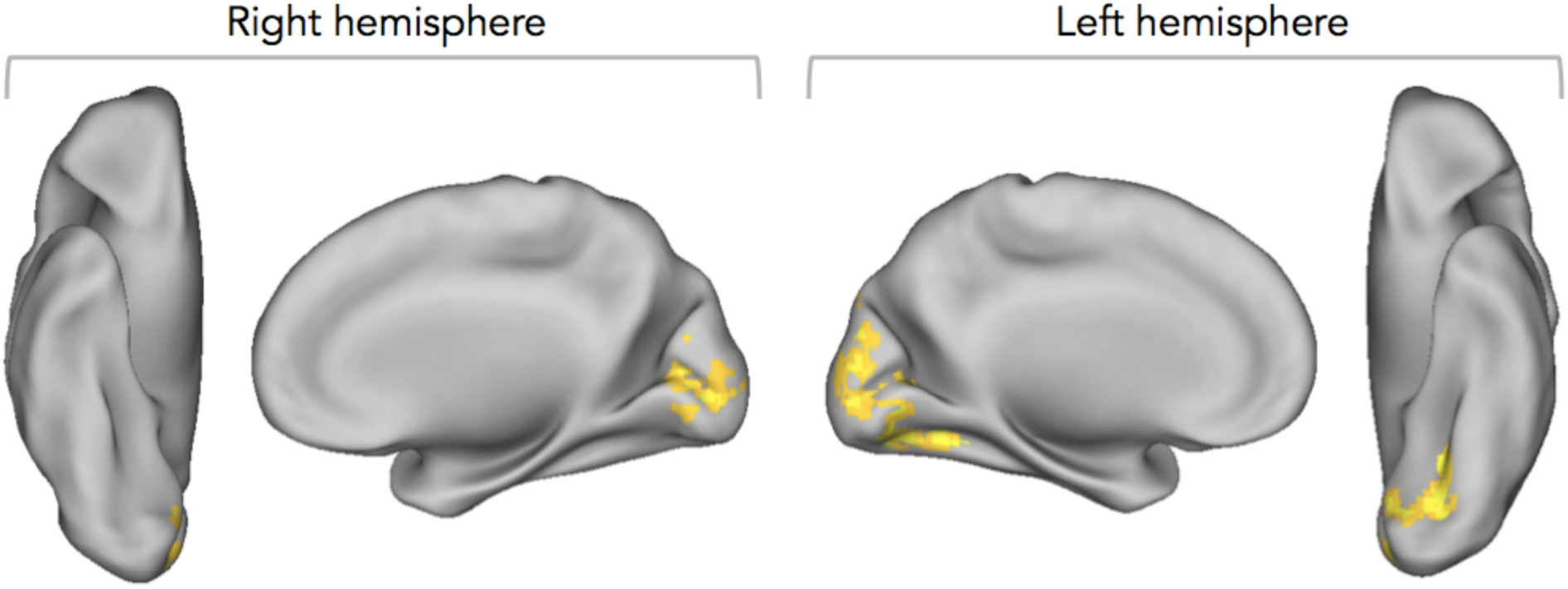
Word-likeness and lexicality effects for fMRI study 1 (Evaluation B). In the main text, we reported an effect of word likeness in occipito-temporal regions posterior to the word-sensitive lvOT cluster in our second fMRI experiment (event-related single trial design; Fig. 3l). While the blocked design of fMRI study 1 was not primarily designed to demonstrate such stimulus-specific effects, we nevertheless also subjected this data set to an event-related analysis of word-likeness. Word-likeness, modeled as a continues factor, produced a more widespread activation effect in fMRI study 1, distributed over occipital regions of left and right hemisphere, with greater activity for more word-like letter strings (two significant clusters: Cluster 1: peak voxel at x = −12, y = −73, z = 1; Left lingual gyrus; T =7.34; 514 voxels; Cluster 2: peak voxel at x = −6, y = −88, z = 37; Left precuneus; T = 4.0; 67 voxels). From the ventral view of the left hemisphere it is visible that the cluster extended into the posterior lvOT, which is not the case in the right hemisphere (Activation effects are visualized at voxel level *p* <. 001 uncorrected; cluster level *p* <. 05 family-wise error corrected). In addition, we tested the words > pseudowords contrast: no significant activation difference between words and pseudowords was found. Only when neglecting the cluster correction, a small activation cluster was found in left frontal cortex (x = −39, y = 38, z = 25; Left frontal pole; T = 3.7; 7 voxel). To summarize, consistent with the second fMRI experiment, an effect of word-likeness on brain activation was found posterior to lvOT, while the (weak) lexicality effect was observed anterior to lvOT, i.e., in downstream regions of the frontal lobe.

**Supplementary figure 5.**
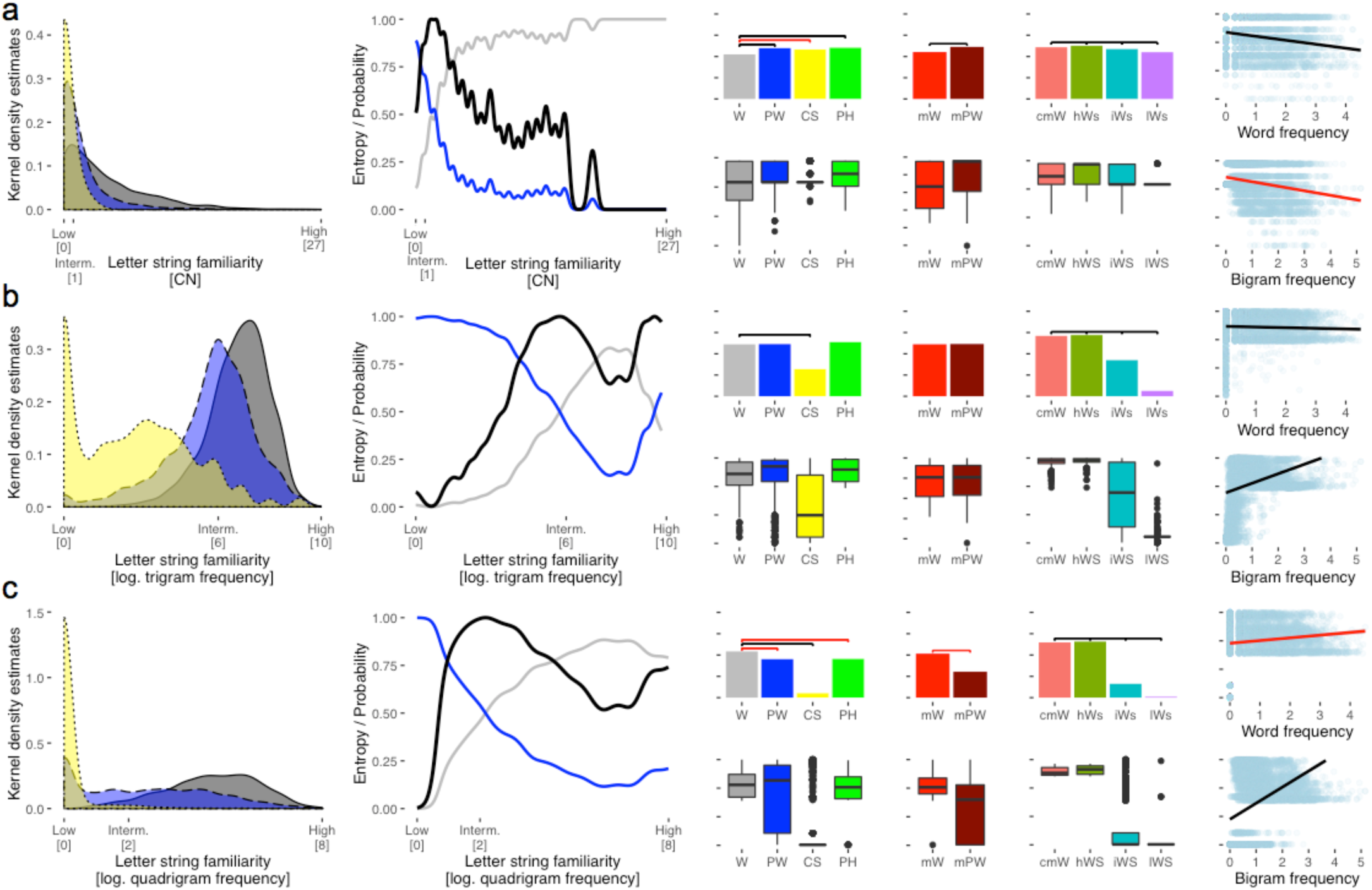
LCM implementations on the basis of three alternative word-likeness measures, plus the simulations of benchmark effects from the literature (Evaluation 1). In the main text, we report an implementation of the LCM using OLD20^25^ as a measure of word likeness that has been reported in the literature to outperform other measures of word likeness REF. However, it is also possible to implement the LCM based on alternative measures of word-likeness. Here, we report three simulations of the benchmark effects tested in Evaluation 1 (cf. Figure 2), using (a) Coltheart’s neighbors, (b) trigram frequency, and (c) quadrigram frequency, as bases for the LCM simulations. The left-most columns show the distributions of the respective word-likeness measure for different types of letter strings as well as the probabilities of being a word or not and the resulting entropy (categorization uncertainty), analogous to Figure 1 in the main text. It is visible that all three measures are less well able to distinguish between words, pseudowords, and consonant strings than OLD20 (Fig. 1a) does. As a result, the resulting entropy function has a different shape than the one derived from OLD20. The LCM implementation based on OLD20 (Fig. 1 and 2) clearly outperformed (in terms of correctly predicted effects and estimated effect sizes) these models based on alternative word-likeness measures. When inspecting the pseudoword > words^13^ contrast, only the model based on Coltheart’s N (Suppl. Fig. 5a) was able to predict this difference; on the other hand, this was the only model that did not predict the contrast words > consonant strings^2^. For description of labels see Figure 1 and 2.

**Supplementary figure 6.**
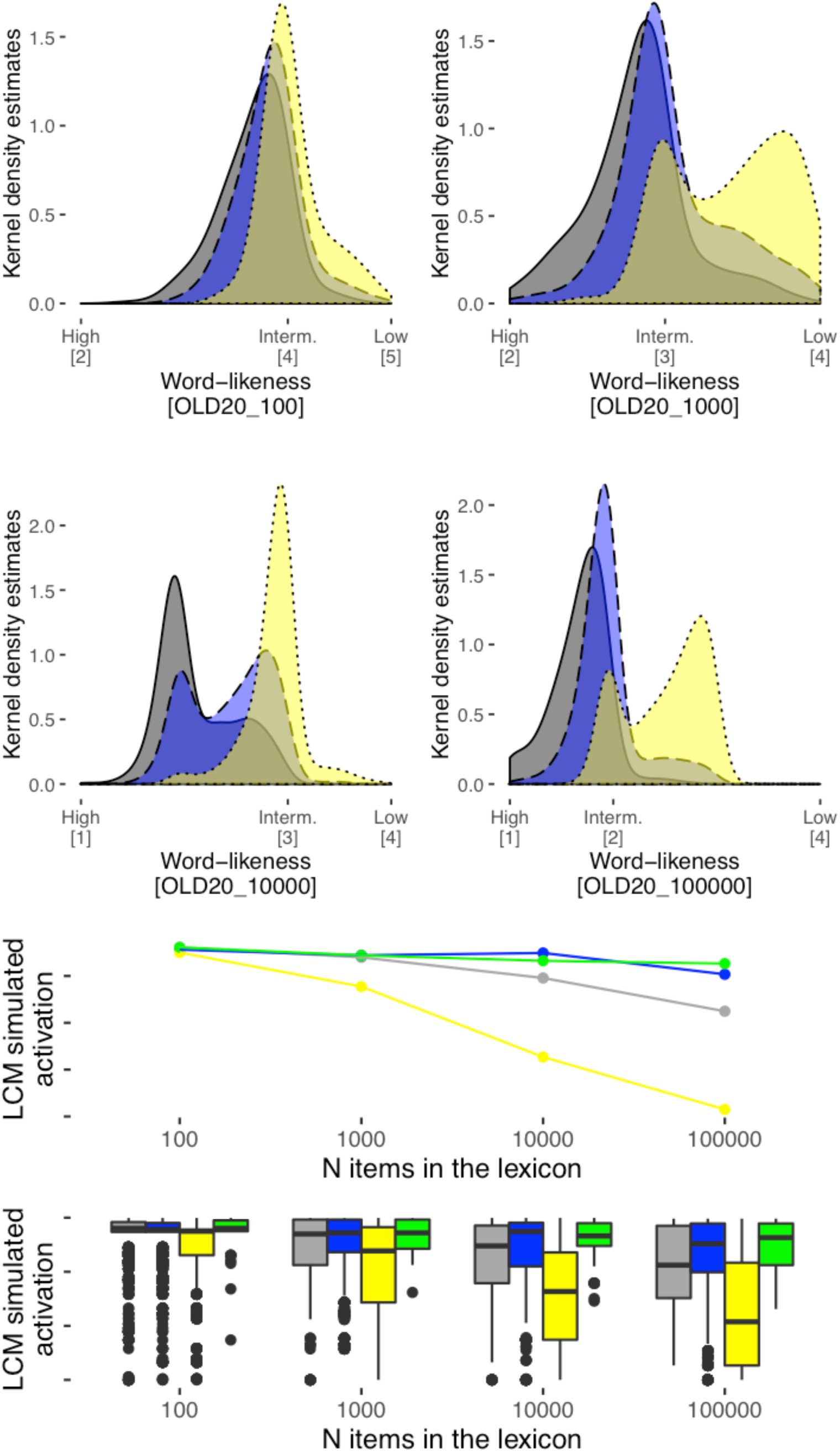
Effect of lexicon size on word-likeness estimations and LCM simulations. We assumed that lexicon size influences word-likeness estimations (i.e. the number of items in the lexicon to which e.g. the OLD20 is estimated) and LCM simulations. First, word-likeness distributions for words, pseudowords, and consonant strings, which were used for the LCM model (see Fig 1 & 2; see Materials section), are presented for lexica consisting of the most frequent 100, 1,000, 10,000 and 100,000 words of the SUBTLEX database. When comparing the word-likeness distributions, it becomes obvious that increasing the size of the lexicon results in a better differentiation between letter string categories (e.g., stronger differentiation between words and consonant strings). Simulations from LCM models derived from these distributions (compare to Figure 1 & Supplement Figure 5), in the lower panels (line graphs show median LCM simulated activation), showed that the model predicted no difference between categories with very small lexicons. Lexicons with intermediate size already allow a differentiation between consonant strings (yellow) and the other stimulus categories. Starting from lexicons with 10,000 words, clearer differentiation between words (gray) and pseudowords (blue)/pseudohomophones (green) was present. In part, besides established effects such as acquired letter knowledge or grapheme to phoneme conversion (for example^13^), these simulations demonstrate that the increasing lexicon size may account for critical patterns of developmental change during literacy acquisition; our present work, in this context, suggests that the lvOT may be an important mediator of such developmental processes.

